# Division of labor in cargo and membrane recognition by SNX1-SNX5: Insights from multiscale modelling

**DOI:** 10.1101/2025.04.27.650847

**Authors:** Satya Chaithanya Duggisetty, Gaurav Kumar, Krishnakanth Baratam, Anand Srivastava

## Abstract

Sorting Nexins (SNXs) are a large group of diverse cellular trafficking proteins that play essential roles in membrane remodeling and cargo sorting between organelles. SNX proteins comprise a banana-shaped BAR domain that acts as a curvature-inducing scaffold and a phosphoinositide lipids sensitive Phox-Homology Domain (PXD) that interacts with the membrane to ensure specific and efficient organelle binding. In concert with the larger retromer machinery, these proteins traffic and recycle cargo between the endosomal membrane, trans-Golgi network, and plasma membrane. Interestingly, the SNX1-SNX5 heterodimeric construct forms a part of the newly discovered pathway where cargo sorting and membrane remodeling can take place in a retromer-independent fashion. In this work, we use molecular dynamics and continuum mechanics simulations to understand the features of SNX1-SNX5 heterodimer, especially the molecular determinants at PXDs, which impart organelle membrane specificity and retromer-independent cargo recognition ability to these proteins. Our all-atom molecular dynamics simulations with isolated PXDs and full-length SNX1-SNX5 on bilayers show that SNX1-PXD has robust membrane-binding features that are largely insensitive to single or double mutation of the basic residues on its surface. Comparing the simulation-based binding poses against the recently solved cryo-EM structures of tubular membrane-bound SNX1 homodimer and SNX1-SNX5 heterodimer also provided interesting insights into the association profile of isolated PXDs when they have the freedom to explore different membrane-binding poses. Our protein-protein simulations of SNX5-PXD with the tail region of the CI-MPR transmembrane cargo protein using metadynamics simulations reveal aromatic residue-rich *π− π* interactions between the two proteins, and a favorable and kinetically accessible binding free energy profile for SNX5. To model the emergent behavior of cargo sequestration and endosomal tube formation by SNX1-SNX5 heterodimer, we also performed Dynamically Triangulated Surface (DTS) based mesoscopic simulations by developing an augmented Helfrich-like continuum-mechanics Hamiltonian to incorporate transmembrane proteins in DTS models.

**SIGNIFICANCE:** Sorting Nexins (SNXs) are a conserved family of endosomal coat proteins that drive cargo-rich tubule formation for receptor transport and recycling within the cell. The cargo-recognition and sorting in these pathways are mediated by the associated retromer complex. SNX1-SNX5 heterodimer is unique to the SNXs family as it has been shown to sequester cargo for trafficking in a retromer-independent manner. The canonical membrane adaptor Phox-homology domain of SNX5 has evolved to carry out this special function. Using molecular dynamics and advanced sampling metadynamics calculations as well as Helfrich-like Hamiltonian-based mesoscopic modelling, we explore the molecular and thermodynamic driving forces that makes the SNX1-SNX5 heterodimeric constructs multifunctional in character.

## INTRODUCTION

Sorting Nexins (SNX) are a large family of multidomain peripheral proteins that predominantly exist as coat proteins in endosomal pathways and regulate organelle cargo sorting, membrane remodeling, trafficking, and signaling processes between the endosomal membrane, plasma membrane, and trans-Golgi network (1–4). Since the balance between protein recycling and degradation is crucial to maintain cellular homeostasis, inappropriate sorting or transport defects often lead to neurological disorders (5, 6). SNX family is ubiquitous and found across phyla, from yeast to mammals, with 33 known mammalian SNXs identified so far \cite{yang2019emerging}. Based on the types of domains in SNX, they are categorized into 5 subfamilies: SNX-BAR (Bin/Amphiphysin/Rvs), SNX Phox Homology Domain (SNX-PXD), SNX-FERM, SNX-PXA-RGS-PXC, and other SNX subfamilies (1). SNX proteins are a crucial component of the retromer machinery, a multi-subunit protein complex responsible for recycling or retrograde trafficking of transmembrane proteins, termed as cargoes. These cargoes include CI-MPR, EGFR, CD44, GPCRsWnt sorting receptor and transporters, among others (2, 7), that are transported between endosomes, the trans-Golgi network (TGN) and the plasma membrane. The mammalian retromer complex comprises a multiprotein subunit of VPS (vacuolar protein sorting associated protein) proteins for cargo recognition and a second subunit of SNX proteins for membrane association. The division of labor for cargo sorting (via VPS) and membrane recognition and remodeling for trafficking (via SNX-BAR-PXD) is quite evident in this retromeric trafficking machinery.

All 33 mammalian SNX family members contain the evolutionarily conserved 100–130 residues long membrane adaptor Phagocytic Oxidase (PHOX) homology domain (PXD), which facilitates the interaction of SNXs with the organelle membrane (8). All the SNX-PXDs in the mammalian family have more than 50 % sequence similarity with SNX1-PXD (except for SNX26) (9) and have largely similar structures except for PXDs of SNX5, SNX6 and SNX32 (see Fig. 1). PXD is considered to be a well-known phosphoinositide-binding domain, and several PXDs are thought to bind with endosomal PI3P lipids, as reported in studies with yeast Vam7, human SNX3, and p40phox (10–12). Most PXDs of SNX family proteins preferentially bind PI3P with few exceptions that interact with other PIP lipids (7) The structures of the PXD of several SNX family and other PXD-containing proteins are characterized previously (8). They share a common core secondary structure comprising of a three-stranded *β*-sheet (*β*1–*β*3) connected by three *α*-helices (*α*1–*α*3) and an irregular strand containing the PXXP region (also known as the PPK loop). The PXD contains two lipid-binding sites: a canonical site for interacting with PIP lipids and a non-canonical site for interacting with PS and PC lipids. Two conserved ARG residues were identified as a part of PI3P binding sites in respresentative PXDs like p40, p47 and SNX3 PXDs. In SNX1-PXD, the corresponding ARG residues are ARG186 and ARG236, which form the canonical PIP binding site. Apart from this, the PPK loop and particularly, the LYS214 also form an important lipid binding site (8, 13). Mutational studies of SNX1 have provided additional insights into the membrane-binding property of the SNX1 PXD. In a liposome study, triple mutations at TYR194ALA, LYS196ALA, and LYS200ALA in the SNX1-PXD were found to significantly affect its membrane-binding ability. However, a single mutation at LYS214ALA does not significantly affect membrane binding even though it is a part of the PPK loop (13). Another mutagenesis study has shown that double mutation of ARG185ALA (ARG186ALA in human) and LYS225ALA (LYS226ALA in human) affects membrane binding and membrane remodeling ability of SNX1 to some extent since they observed reduction in membrane tubulation. By adding a third mutation at PHE186ALA (PHE187ALA in human), membrane tubulation is also prevented (14). These biochemical assays clearly suggest that the membrane-binding ability of SNX1-PXD is a coperative in nature and results from a combined effect of several residues distributed on the structure.

**Figure 1:**
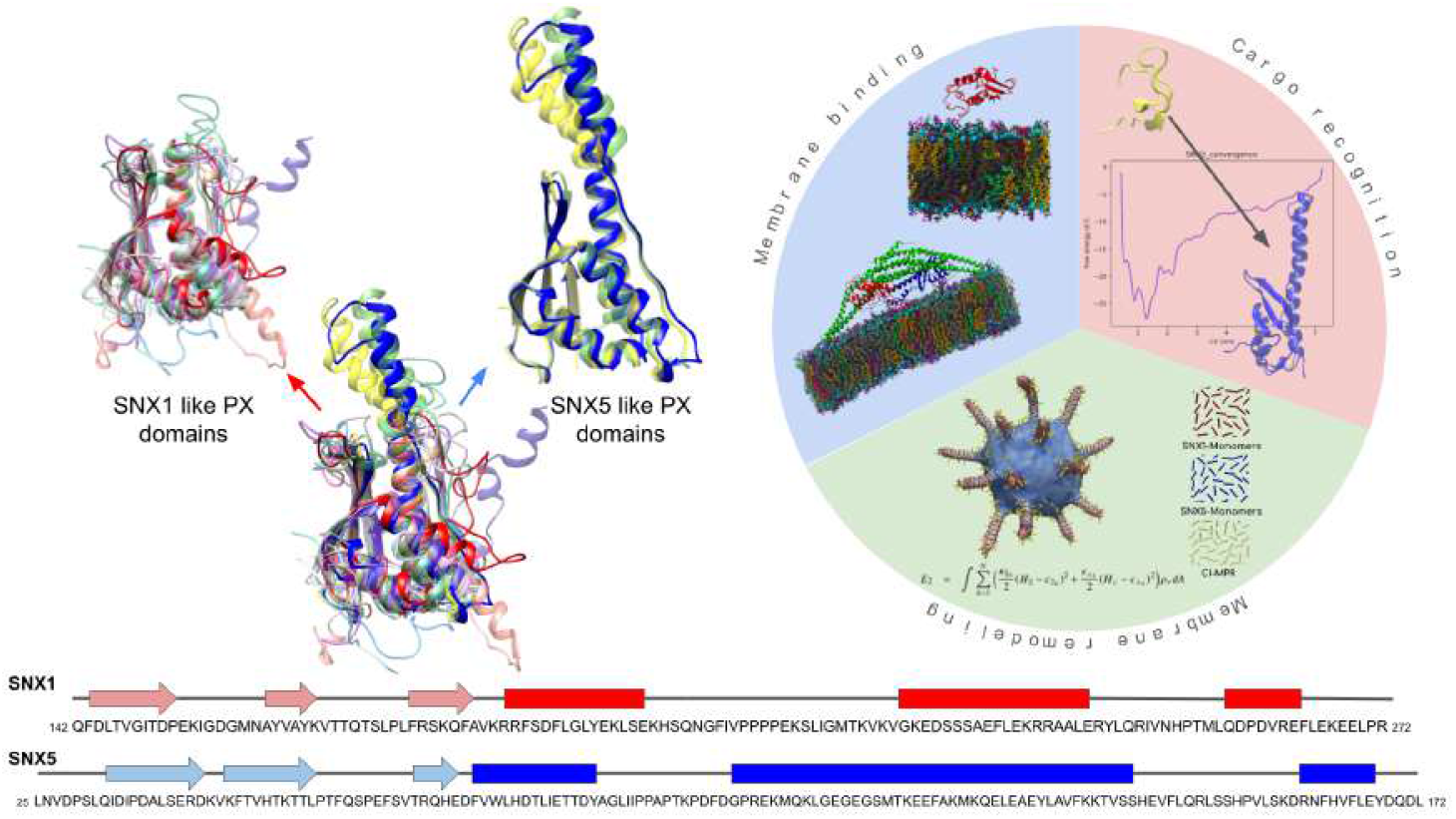
Structure and Functional properties of SNX. The similarity in the structures of SNX proteins is depicted by superposing 16 known structures of different SNX proteins. The SNX5 (SNX5, SNX6, and SNX32))-like proteins can be clearly distinguished from SNX1-like proteins with their long *α*2 helix (labeled in the left figure). The sequence and secondary structure information of SNX1 (red) and SNX5 (blue) are also shown. In this study, we have addressed the membrane binding, cargo recognition and membrane remodeling properties of SNX1 and SNX5 using computational approaches at various resolutions

Though SNX-PXDs are primarily lipid-binding domains, data from Peter Cullen’s group at University of Bristol showed that some cargo recognition can also take place via special motifs present in the PXD of SNXs (SNX5/6 in this case) (15–18). This mechanism is now commonly known as “endosomal SNX-BAR sorting complex for promoting exit-1 (ESCPE-1)”. This transformative discovery establishing that ESCPE-1 and retromer are functionally distinct and that the specific recognition of cargo was carried out by PXD of a SNX expands the role of PXD beyond membrane adaptor. Through X-Ray crystallography and biochemical studies, Cullen group showed that SNX5-PXD carried a very unique sequence motif that specifically recognizes and binds to a beta hairpin like structural motif in the transmembrane cation-independent mannose 6-phosphate receptor (CI-MPR) cargo (18). Notably, there are over 60 other cargo proteins that are known to have such a beta hairpin motif and require SNX5/6 for their recycling. Interestingly, SNX5/6 does not exist as homodimer but forms functional heterodimer with SNX1/2, which can now carry out the dual role of cargo recognition as well as the BAR-domain driven formation of tubulo-vesicular transport carriers (19).

Amino-acid sequence in PXD of SNX5/6 is unique since both SNX5 and SNX6 PXDs lack the two conserved ARG residues involved in PI3P binding. Also, it has an additional 15-residue insertion immediately after the PXXP motif (20) which increases the length of *α*2 helix as shown in Fig.1. Interestingly, the PXXP motif is expanded into a double PXXP motif with the sequence PXXPXXP. The differences in sequence and structure between the two domains are clearly shown in the left panel of Fig.1. In Fig. S1, we show the differences in their electrostatic profiles. SNX5-PXD has very few basic residues on the surface, which significantly affects their anionic lipid recognition. Though SNX5/6-PX domains bind weakly with different PIP lipids, they play an important role in retromer function. Changes in length and sequence of SNX5/6-PXD profoundly impact the specific structure and conformation required for its cognate membrane partner recognition. The cargo recognition happens via a motif on SNX5/6 that is missing in SNX1/2. To perform its function, SNX5/6 forms heterodimers with SNX1/SNX2 and binds to transmembrane proteins from endosomal membrane for transport to TGN or plasma membrane (2). SNX1/2 binds to the PI3P /PI(3,5)P2 and induces curvature on endosomal membrane leading to tubulo-vesicular formation. The cargo CI-MPR binding with SNX5-SNX1 heterodimer has been established using GFP tagging experiment, where the residues 2347-2375 of CI-MPR tail were identified as essential for the SNX5/6 association (18).

The structure of SNX5-PXD bound to CI-MPR tail region was determined previously using X-ray crystallography method (PDB IDs: 6N5X, 6N5Y) (21). The structural data shows that CI-MPR peptide interacts with the SNX5-PXD domain via a *β*-sheet augmentation, forming two anti-parallel *β*-strands (*β*A and *β*B) connected by a long flexible linker. The two structures vary in the amino acids forming the second beta-strand, causing a flip in the side chains and interactions with SNX5. Due to this, PHE136 interacts with LEU2370 in 6N5X and MET2371 in 6N5Y. Moreover, an Aspartate mutagenesis study showed that ^2349^*V SY KY SK*^2355^ region in CI-MPR is important for SNX5 binding(21). Also, the recent 10 Å resolution membrane-bound cryo-EM structures of SNX1 homodimer (14) and SNX1-SNX5 heterodimer (22) have shed unprecedented light on the membrane association profile of SNX and also on the protein-protein interactions leading to large coat formation on the tube. In the presence of the BAR domain and the amphipathic helix (AH), the primary membrane-binding components of SNX1 (14), mutation at the canonical PXD lipid binding site showed no significant effect on membrane binding. Several mutations on SNX1-PXD were tested for its effect on membrane association, and only a few cases showed any effect on membrane binding and membrane remodeling. This suggests that SNX1 may possess multiple lipid-binding sites dispersed across its surface with varying binding affinities. Similarly, CI-MPR explores more than one stable conformation with SNX5-PXD, which might indicate the adaptability of SNX5 to bind various cargo proteins. In our study, we explore the kinetic accessibility of SNX PXDs with their interacting partners using all-atom molecular dynamic simulations. In this work, we use all-atom molecular simulations on PXD-Bilayer systems, full-length SNX1-SNX5-Bilayer system, and advanced sampling metadynamics simulations of PXD-bound CI-MPR motif to understand the molecular thermodynamics processes through which the membrane and cargo recognition take place in this retromer-independent cargo-sorting and trafficking heterodimeric complex (see Fig.1). Additionally, we develop and use mesoscopic Helfrich-like Hamiltonian-based continuum mechanics simulations to understand the implications of these molecular interactions on emergent behavior, such as large-scale cargo sorting, tubulation, and trafficking. We juxtapose our atomistic and mesoscopic ensemble trajectory data against the published structural data from cryoEM and x-ray crystallography, as well as against the biochemical and cell biological data to gain additional insights into the molecular driving forces leading to cargo sorting and tubulated coat formation.

The remainder of this article is structured as follows. This Introduction section is followed by the Materials and Methods section, where we list the details of various all-atom systems considered for simulations in this work, along with the parameters used to run these canonical MD and Metadynamics simulations. We also provide details of our continuum mechanics Dynamic Triangulated Surface (DTS) model (23–28) to study the larger scale membrane deformation. Towards that, we have enhanced our DTS Hamiltonian formulation to allow for the sorting of transmembrane cargoes, which we discuss in detail in this section. The Material and Methods section is followed by the Results and Discussion section, where we report our findings. We show the ability of SNX1 to bind different membrane compositions and the role of aromatic residues in SNX5 to bind various cargo proteins. With the mesoscopic continuum modeling, we demonstrate the impact of homo or heterodimerization of SNX5 and SNX1 on membrane remodeling and cargo transportation. We summarise our findings in the Conclusion section. All input files needed to initiate molecular simulations and mesoscopic continuum mechanics simulations, as well the full trajectory data of all simulations for all systems considered in this work, are publicly available on the following Zenodo repository:https://shorturl.at/Id7FI. We have also deposited extended versions of the movie files that are part of the Supporting Information (SI) in the same repository.

## MATERIALS AND METHODS

### 0.1 All-atom molecular dynamics simulations

We performed a series of all-atom molecular dynamics (AAMD) simulations of several protein-bilayer systems (listed in Table 1). We chose three different lipid bilayer systems that represent the endosomal membrane (POPC: POPS: POPE: PI3P: PI35P:: 50:15:25:8:2), plasma membrane (POPC: POPS: PI45P:: 80:19:1) and Golgi membrane (POPC:POPE:PSM:POPS:PI4P:CHL:: 50:17:12:4:9:8). The structures for isolated SNX1-PXD and SNX5-PXD were obtained from Protein Data Bank IDs 2I4K (29) and 6N5X (18), respectively. The structure of full-length SNX1 homodimer was taken from the recently solved membrane-bound cryoEM structure (PDB ID: 7d6d (14)), while the SNX5 was modeled using Alphafold. SNX5 was augmented over SNX1 monomer to make a full length SNX1-SNX5 heterodimer. For the full-length system, the membrane-binding orientation from the solved structure was used to model the initial membrane-bound state. For the solution-state simulations of isolated SNX1-PXD and SNX5-PXD, we positioned the protein away from the membrane such that different membrane binding profiles could be explored. Moreover, we also carried out replica exchange simulations of isolated PXDs on membranes to establish the convergence of binding profiles. We used the recently developed replica exchange with hybrid tempering for peripheral membrane proteins (REHT-PMP) for this purpose (30, 31). The details of the REHT-PMP simulations are discussed in the next section.

**Table 1:** Details of atomistic molecular simulations performed in this study.

| Protein | Membrane | System size (no. of atoms) | Trajectory length (μs) |
| --- | --- | --- | --- |
| SNX1 | Endosomal | 72227 | 2.0 |
|  | Plasma | 74522 | 1.0 |
|  | Golgi | 76846 | 0.5 |
| SNX5 | Endosomal | 101104 | 2.0 |
|  | Plasma | 75772 | 1.5 |
|  | Golgi | 74413 | 0.5 |
| SNX1-SNX5<sub>FL</sub> | Endosomal | 596453 | 2.0 |

**Table 2:** Details of all-atom REHT simulations performed in this study.

| Protein | Membrane | System size (no. of atoms) | no. of replicas | Trajectory length (μs) |
| --- | --- | --- | --- | --- |
| SNX1 | Endosomal | 86614 | 15 | 1.5 |
|  | Plasma | 74522 | 15 | 1.5 |
|  | Golgi | 76846 | 15 | 1.5 |
| SNX5 | Endosomal | 101104 | 20 | 2.0 |
|  | Plasma | 75772 | 20 | 2.0 |
|  | Golgi | 74413 | 15 | 1.5 |

CHARMM-GUI (32) was used to set up initial systems and we have used the CHARMM36m force field for our molecular dynamics simulations. The modeled systems were energy minimized and equilibrated for 20 ns and a combination of NVT/NPT simulations were carried out for initial stabilization of the bilayer-protein systems. This was followed by production runs where each system was simulated at a temperature of 310 K in the NPT ensemble using GROMACS 2021.3. (33). The temperature was controlled using Nose-Hoover thermostat (34, 35) and semi-isotropic pressure coupling to set the pressure to 1 bar was achieved using Parrinello-Rahman barostat. (36). The duration of the production run for each system is reported in Table I. The trajectories were stored at a dump frequency of 1 ps.

For the isolated PXD-bilayer system, a bilayer patch of about 200 lipids was set up with roughly 23,000 TIP3 water molecules and 150 mM NaCl salt concentration. For the larger full-length SNX1-SNX5 dimer on the membrane, 1200 lipid molecules and 141,536 TIP3 water molecules with 150 mM NaCl concentration were used. The system size details are also provided in Table 1. The z-analysis was carried out using an in-house VMD TCL script and Python code was used for plotting. lipid wise contact analysis was done using gmx hbonds and a python script was used to visualize.

### 0.2 Replica Exchange with Hybrid Tempering for Peripheral Membrane Proteins (REHT-PMP) simulations

The initial configurations for setting up aa Replica Exchange with Hybrid Tempering for Peripheral Membrane Proteins (REHT-PMP) simulations were obtained by dumping the membrane-bound structure from the six systems described in Table1 and were run using GROMACS 2022.5 (37) patched with PLUMED 2.9.3 (38). A total of 15 or 20 replicas were used based on the exchange probabilities 2. The interactions in these systems were defined by the CHARMM36m force field (39) and TIP3P water model (40). The membrane and solvent experience temperatures between 310 and 330 K across the replicas. However, the protein is allowed to experience both temperature and Hamiltonian scaling from 310 to 430 K. The production run was performed using a timestep of 2 fs in the NPT ensemble. All the non-bonded interactions within 12 Å were evaluated (with a 10 Å switching distance), while the long-range electrostatic interactions were dealt with using the Particle Mesh Ewald method. The Nose–Hoover thermostat and Parrinello-Rahman (with semi-isotropic coupling) barostat with time constants 1.0 and 5.0 *ps*^*−*1^ respectively, were used to maintain the defined temperature of the replica and to maintain a pressure of 1 atm. All the individual replicas were equilibrated for 1 *ns* at their respective temperatures before the exchanges among replicas were initiated. The exchange was attempted every 1 *ps*. Each of the replicas were evolved for a total of 100 ns, thus leading to a total simulation time of 1.5 or 2.0 *μs*. Further details of this method, along with the relevant codes and setup files can be found in our previous work (41).

### 0.3 Cryo-EM analysis

We used the coordinates of the full-length SNX1-SNX5 construct (PDB: 8AFZ) deposited along with the 10 Å cryo-electron density map (EMD-15413) for our full-length SNX1-SNX5 simulations on the bilayer (22). However, since our isolated PXDs gave us degenerate binding profiles, we wanted to see if the same electron density could tolerate the alternative binding poses that we were getting from AAMD simulations. To model the heterodimer with this alternate SNX1-PXD orientation, we included the BAR domains coordinates of SNX1 from the PDB ID: 8AFZ and modeled the amphipathic helix that connects this BAR domain with our alternate SNX1-PXD using I-TASSER (42, 43). This heterodimer with alternate SNX1-PXD orientation was fit into the cryo-ET map using iMODFIT (44). The protein coordinates and the electron density map were visualized using UCSF-chimera (45). For visualization of the fitting process, please see the movie file movieS1.mpeg in SI.

### 0.4 Protein–ligand Metadynamics simulations

The Coordinates of SNX5 with CI-MPR tail region (PDB: 6N5X) were used to generate the initial configuration files for the setup of the metadynamics simulation. On the other hand, SNX1 with CI-MPR was set up by structurally aligning SNX1 with SNX5, preserving the orientation of CI-MPR as observed in the above PDB file. Each of these systems were then solvated with TIP3P water molecules and neutralized using counterions equivalent to 150 mM of NaCl. All the systems were energy-minimized and equilibrated to a temperature of 300 K and 1 atm pressure before proceeding to production runs. Temperature and pressure of the simulation systems were maintained by using Velocity rescaling algorithm (time constant of 0.1 ps) and Parrinello–Rahman algorithm, respectively. LINCS algorithm was used to constrain all bonds involving hydrogen atoms. The long-range electrostatic interactions were treated with particle mesh Ewald scheme, while the Verlet scheme was used to treat the short-range interactions with a cutoff of 1.4 nm. All the simulations were performed in GROMACS patched with PLUMED-2.4.8. The collective variable(CV) was designed based on the available experimental information about the interaction of PHE136 of SNX5 and LEU2370 of CI-MPR. Hence, the distance between the alpha carbons of these two residues was used as the CV of interest to obtain the free energy landscape of binding of SNX5 with CI-MPR. Similarly, in the case of SNX1, upon structural alignment with experimentally resolved SNX5-CIMPR complex, the equivalent residue of interaction was found to be LEU236 of SNX1 and LEU2370 of CI-MPR. Hence, we used the distance between the alpha carbons of these two residues as the CV. The interactions and contacts between SNX-PXDs and CI-MPR were extracted using the pyContacts tool ((46)).

### 0.5 Mesoscopic Hamiltonian: Membrane as dynamically triangulated continuum surface with SNX and TM cargo proteins as anisotropic inclusions

In the classical mesoscopic DTS model, the fluid membrane is modeled as a dynamically triangulated surface and proteins are modeled as nematics. These nematics 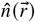 lie on the vertices of the triangulated membrane and are free to move in the local tangent plane of the associated vertex (23, 26). An extended Helfrich-like Hamiltonian is solved to arrive at the remodeled membrane morphology due to combination of intrinsic properties of individual components (such as bending modulus of membrane and curvature of the proteins) and interactions between them (such as binding energies between membrane and proteins and protein-protein interactions). Recently, we developed an advanced version of this continuum-scale DTS model such that multiple types of curvature-inducing proteins can interact with each other on the membrane surface (27). The augmented Hamiltonian allowed us to study the emergent membrane remodeling effects when different curvature-inducing proteins were allowed to interact with each other on the membrane surface. In the previous formulation, the proteins were free to move on the surface of membrane but they didn’t have the ability to share the same node on the triangulated surface, which was addressed in our previous study (27). This expanded the scope to study systems such as retromer complex where multiple proteins exist as an oligomeric unit. However, the previous model also did not allow out of plane protein embedding, which would be needed to incorporate transmembrane cargo proteins such as CI-MPR, which is of interest to us in this study. In the current extended version of the DTS model, we have designed the Hamiltonian such that the two different types of in-plane proteins (SNX1 and SNX5) are free to move on the membrane and there is a third protein (CI-MPR) that has the ability of superimposing on the SNX1 and SNX5 proteins in the out of plane manner. We have modeled the CI-MPR as a nematic that is always perpendicular to the vertex tangent plane.

In this model, membrane-protein interaction is modelled as anisotropic spontaneous curvatures of the membrane in the vicinity of the protein filament. Protein-protein interactions are modelled by the splay and bending terms of the Frank’s free energy for nematic liquid crystals. The total energy is the sum of membrane energy, interaction energy between protein and membrane and the protein-protein interaction energy. The total energy is given as: *E*_*T*_ = *E*_*Memb*_ + *E*_*Memb−prot*_ +*E*_*prot−prot*_, where *E*_*Memb*_ is the Canham-Helfrich elastic energy for membranes, (47, 48) *E*_*Memb−prot*_ is the interaction energy between proteins and membrane and *E*_*prot−prot*_ is the protein-protein interactions energy. The membrane energy is given as, 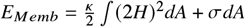, where *E*_*Memb*_ is the Canham-Helfrich elastic energy for membranes, (47, 48) *ϰ* is membrane’s bending rigidity and *H* is membrane mean curvature and *σ* is the surface tension of the membrane. The mean curvature is given as *H* = (*c*_1_ + *c*_2_) / 2 where *c*_1_ and *c*_2_ are the local principal curvatures on the membrane surface along the orthogonal principal directions.

The membrane and protein interaction term in the Hamiltonian is given as: 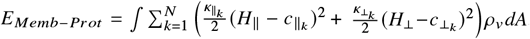, where *N* is the number of protein monomers. In this case, we have two different protein monomers (SNX-1 *&* 5) so we have used *N* = 2. *ϰ*_∥_ and *ϰ*_⊥_ are the induced membrane bending rigidities and *c*_∥_ and *c*_⊥_ are the intrinsic curvatures of proteins along 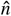 and its perpendicular direction 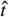 in the local tangent plane. *H*_∥_ and *H*_⊥_ are the membrane curvatures in the direction of 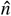 and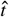, respectively where *H*_∥_ = *c*_1_ cos^2^ *ϕ + c*_2_ sin^2^ *ϕ* and *H*_⊥_ = *c*_1_ sin^2^ *ϕ + c*_2_ cos^2^ *ϕ. ϕ* is the angle between the direction of nematic orientation 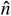 and the principal direction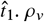 is the local coverage of nematic on the chosen vertex. If a nematic is present on a vertex, *ρ*_*v*_ = 1, otherwise it will be 0. This scheme allows us to have variable surface density of the protein monomers.

The standard protein-protein interaction is derived from Frank’s free energy for nematic liquid crystals and wriiten as: 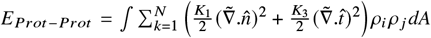 where *K*_1_ and *K*_3_ are the splay and bending elastic constants for the in-plane nematic interactions. 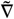 is the covariant derivative on the curved surface. We are using only splay and bending terms for nematic-nematic interaction. For computational purposes, the discrete form of the energy is taken from the Lebwohl-Lasher model (49, 50) and expressed as 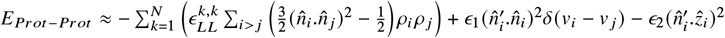,where *ϵ*_*LL*_ is the strength of the nematic interaction with a constant approximation (*K*_1_ = *K*_3_). The sum ∑_*i> j*_ is on all the nearest neighbours *i, j* vertices on the triangulated grid that promotes alignment among the neighbouring orientation vectors. *ϵ*_1_, is the strength of the interaction between out-of-plane and in-plane nematics here (modeling the interactions between the TM cargoes and Sorting Nexins). This term provides the perpendicular interaction between the corresponding nematics (51). Here *v*_*i*_ and *v*_*j*_ are the two neighbouring vertices where the TM Cargo and Sorting Nexin localize and provides the interaction between the two components. The third term provides the perpendicular arrangement of TM cargo molecule (CIMPR in our case) on the membrane. Here, 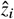 is the normal of the *i*^*th*^ vertex and *ϵ*_2_ is the strength of this term. We use Monte Carlo (MC) moves to evolve the Hamiltonian to the thermodynamic lmits and details of the moves can be found in previous publications (23, 27, 52). Please note that the Monte Carlo moves for the new formulation, which allows overlap of nodes between two nematics, needs to be handled differently than before and we have discussed this below.

### 0.6 Dynamical triangulation Monte Carlo simulations steps

For the Monte-Carlo (MC) simulations to solve the Hamiltonian, we start with a spherical vesicle where all the three nematics are randomly distributed over the surface. As we are using multiple nematics with different orientations, the MC moves are extended as compared to the existing formulations (27, 49, 53). The MC moves have the following sequence: (i) vertex move, (ii) link flips, (iii) In-Plane Nematic (SNXs) shifts, and (iv) In-Plane Nematic rotations (v) Out-of-Plane Nematics (CIMPR)-shifts. The out-of-plane nematics are perpendicular to the membrane local tangent plane at the vertex. These MC moves help the system to reach in the minimum energy configuration and provide the shape relaxation and fluidity in the system.

**Figure 2:**
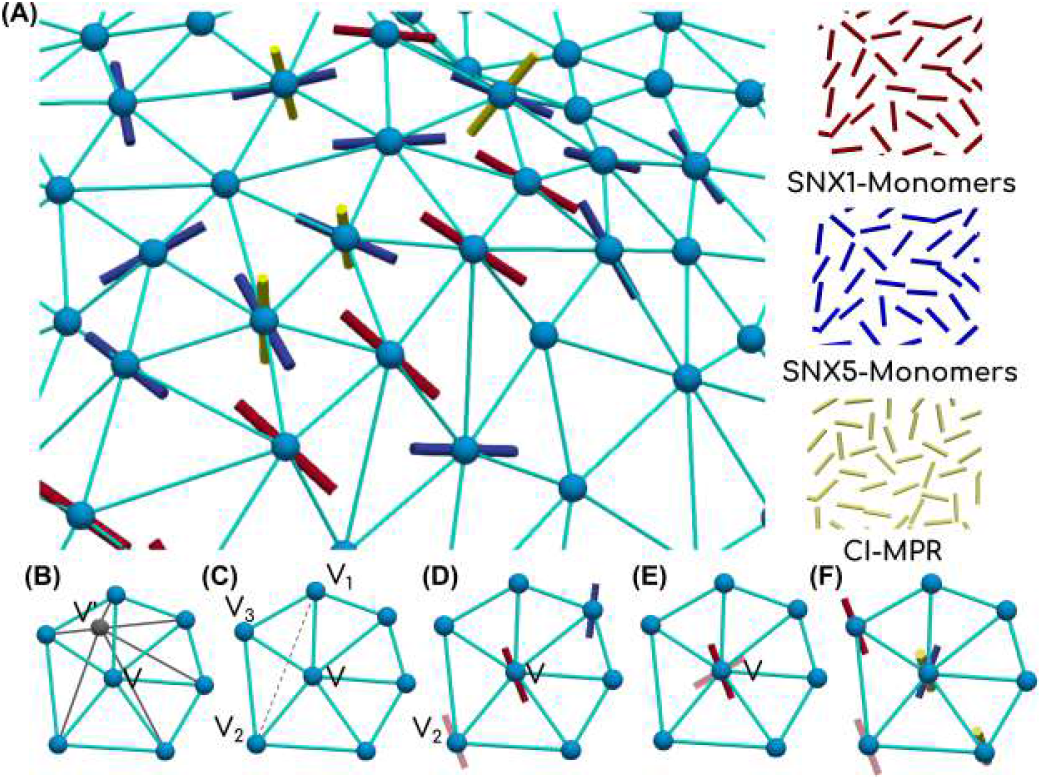
Schematics for the DTS model. (A) The triangulated mesh represents the elastic membrane sheet and the vectors on the nodes represent the proteins as nematics. The model can account for multiple types of nematics presented as different colors in the schematics. (B-F) show the MC moves required for the energy minimization to evolve the shape of triangulated surfaces.

We use Fig.2 to show the new DTS representation. For this work, the Red and Blue nematics in Fig.2 represent in-plane SNX1 and SNX5, respectively. The out-of-plane Yellow nematics represent the CI-MPR cargo protein. The first MC move is shown in the Fig.2 (B) where a randomly chosen vertex is displaced inside the cubical box. In this representation, the box is centered at the chosen vertex, which is displaced from *V* to *V* ^′^. The move explores possible minimum energy configuration for the Neamtic-Membrane system with their interactions. The second MC move is shown in the Fig.2 (C) and called link flip. In this move, a set of two triangles is randomly chosen and the common side between them is removed and a new connection is established between the two disconnected vertices. In the Fig.2 (C), the tether between *V* and *V*_1_ is replaced by creating new link between *V*_2_ and *V*_3_. This move provides the fluidity in the system. The link flip move is known as the dynamically triangulated Monte Carlo (DTMC) move. The third MC move is shown in the Fig.2 (D) that shifts the nematic from one vertex to another vertex. The out-of-plane nematic is modeled in such a way that it will lie in the local normal of the vertex and free to move without any restrictions unlike the in-plane nematics. The fourth MC move allows the nematic rotation in the local tangent plane of the vertex and is applicable for in-plane nematics as shown in Fig.2 (E). This move provides the diffusion of proteins on the membrane surface as it affects protein assembly patterns and drives the membrane towards the minimum energy configuration. The fifth and final MC move is the replica of the fourth move. Forth move works for the in-plane nematics (self- and cross-interactions between SNX1 and SNX-5 here) while the fifth move works for the perpendicular interactions between out-of-plane and in-plane (CIMPR SNX1/SNX5 here) when they both lie on the same vertex and shown in the Fig.2(F).

## RESULTS AND DISCUSSION

### SNX1-PXD exhibits degenerate membrane binding poses with endosomal and plasma membranes

SNX proteins are involved in the endosomal recycling pathway which requires these proteins to bind to endosomal, plasma and golgi membranes. Hence, in our study, we have modeled and simulated the interaction of SNX1-PXD and SNX5-PXD with the three different membrane compositions to mimic the three organelles (Fig. 3). We also simulated the full length SNX1-SNX5 heterodimer on endosomal membrane and compared our data with the available low-resolution cryo-EM data ((22)) (Fig. 4). A detailed description of the membrane composition and the simulation setup is given in the Methods section.

**Figure 3:**
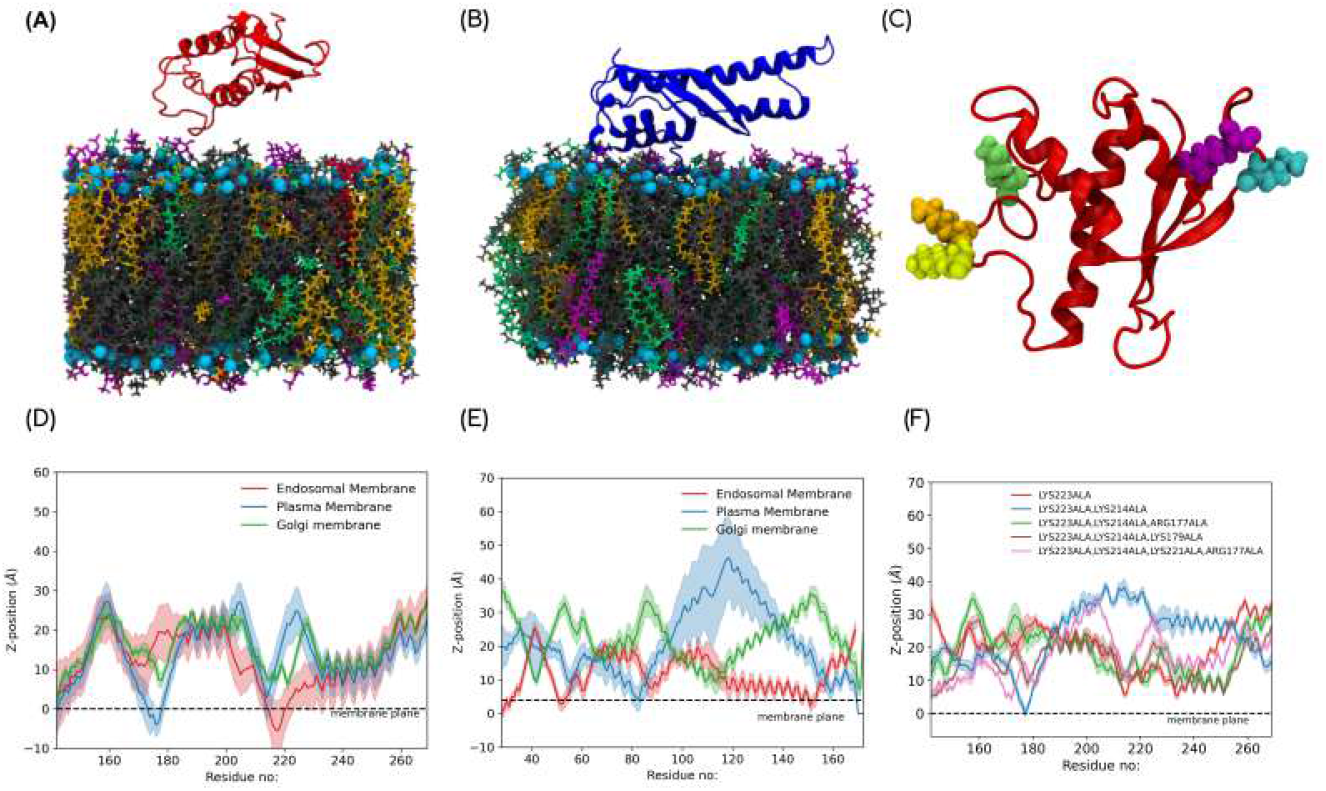
Dynamics of SNX1 and SNX5 PXDs. Illustration of (A) SNX1 (Red) and (B) SNX5 (Blue) PXDs on endosomal membrane with PC, PE, PS, PI3P, and PI(3,5)P represented as grey, orange, green, magenta, and red, respectively. (C) Position of the mutated residues, ARG177, LYS179, LYS214, LYS221 and LYS223 depicted as CPK model in cyan, purple, green, orange, and yellow, respectively. (D, E) shows the z-distance between membrane and PXDs in different membrane compositions, where the x-axis represents the residue number and the y-axis represents the z distance. (F) shows the variation in the z-distance as a result of mutations in SNX1-PXD with endosomal membrane.

**Figure 4:**
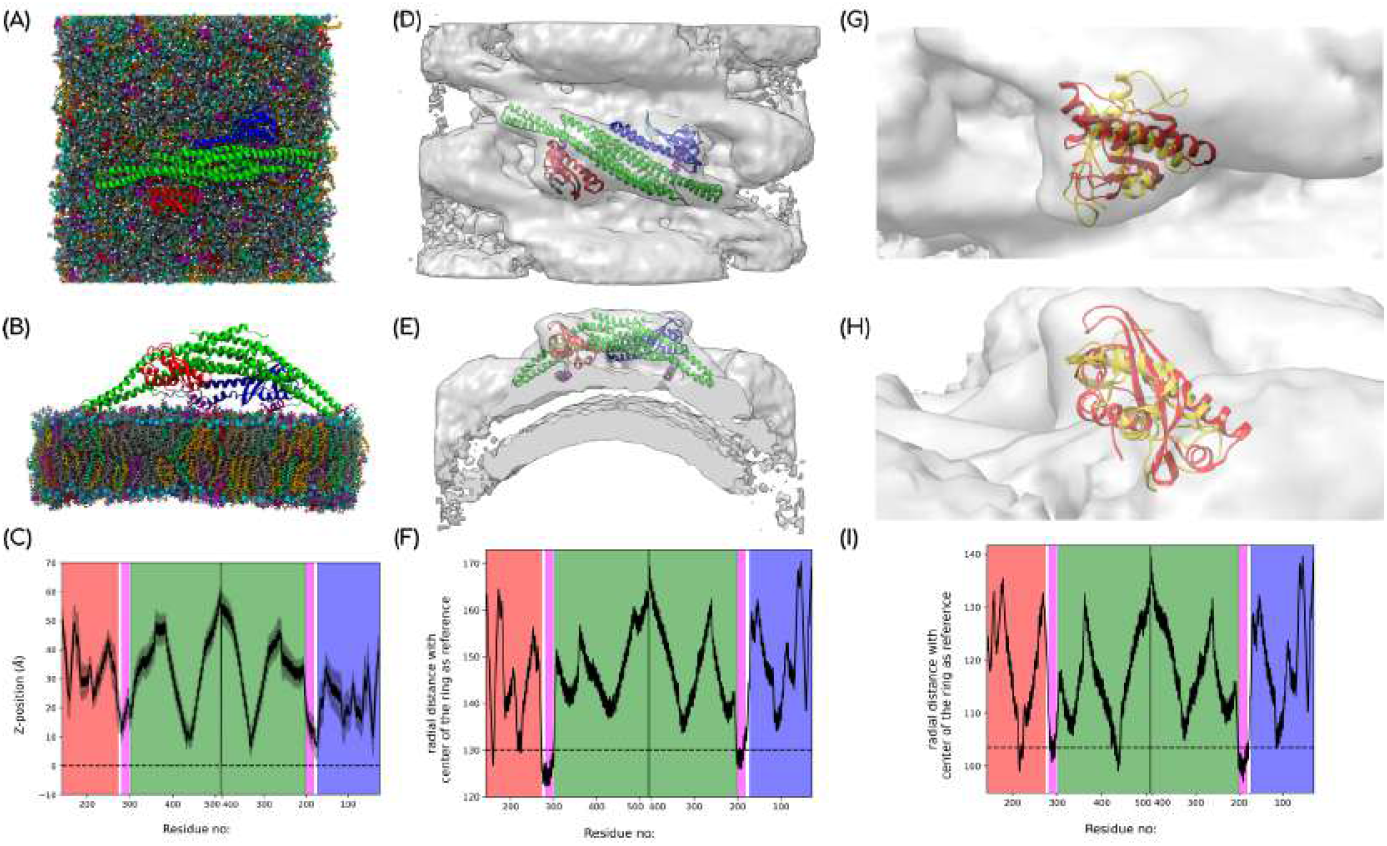
In this illustration, SNX1-PXD/SNX5-PXD, amphipathic helix, and BAR domains are shown in red/blue, magenta, and green, respectively. (A, B) shows the membrane bound SNX1-SNX5 heterodimer on endosomal membrane after 2*s* MD simulation. (D, E) SNX1-SNX5 (PDB ID: 8AFZ) heterodimer fit in the cryo-EM map (EMDB-15413). (G, H) fitting of the SNX1-PXD orientation obtained from our MD simulation (gold) with the cryo-EM map of heterodimer (PDB ID: 8AFZ, red) as discussed in methods0.3. (C, F, I) are the z analysis distances calculated for the respective proteins.

SNX1-PXD contains several positively charged residues that can interact with the negatively charged lipids in the membranes. We calculated the surface electrostatic potential of the SNX1-PXD to understand the extent of the positive charge distribution. Fig. S1 in SI shows the presence of positively charged regions at the loops connecting the three helices along with the surface of *α*3 and the *β*2 - *β*3 loop. In contrast, the solvent-exposed surface of *α*1, *β*3, and the loop connecting the amphipathic helix with *α*3 are negatively charged. In our study, the simulations were started by placing the protein about 10 Å away from the membrane in order to prevent any starting configuration bias. During simulation, the SNX1-PXD reorients and interacts with the membrane and we analyze these unbiased trajectories to gain insights into their membrane binding property. To understand the spatial orientation of the PXD relative to the membrane, we performed Z-distance analysis. This approach involves measuring the distance between the phosphate headgroup plane of the lipid bilayer and the C*α* atoms of each of the protein residues. For the endosomal membrane system, Fig.3(D) shows that the *α*1–*α*2 loop inserts into the membrane. Particularly, the residues LYS221 and LYS223 interact with PE lipids and LYS238 interacts with PIP lipids (Fig.5). In the case of plasma membrane, Fig.3(D) clearly shows the insertion of *β*2-*β*3 loop and *α*2 helix into the membrane. The residues ARG177 and LYS239 bind with POPS lipids and a small percent of PI(4,5)P lipids interact with LYS179. Interestingly, the insertion of PXD in the Golgi membrane is highly similar to the plasma membrane (Fig.5). In all three membranes, the residues in the PPK loop also interact with the POPC lipids. Altogether, the endosomal interaction appears to center around the conserved residue ARG236, in agreement with previous literature. In contrast, SNX1 binding to plasma and Golgi membranes seems to involve regions that were not identified previously. These findings suggest that SNX1 has the ability to bind to different membrane compositions by adopting different binding interfaces.

**Figure 5:**
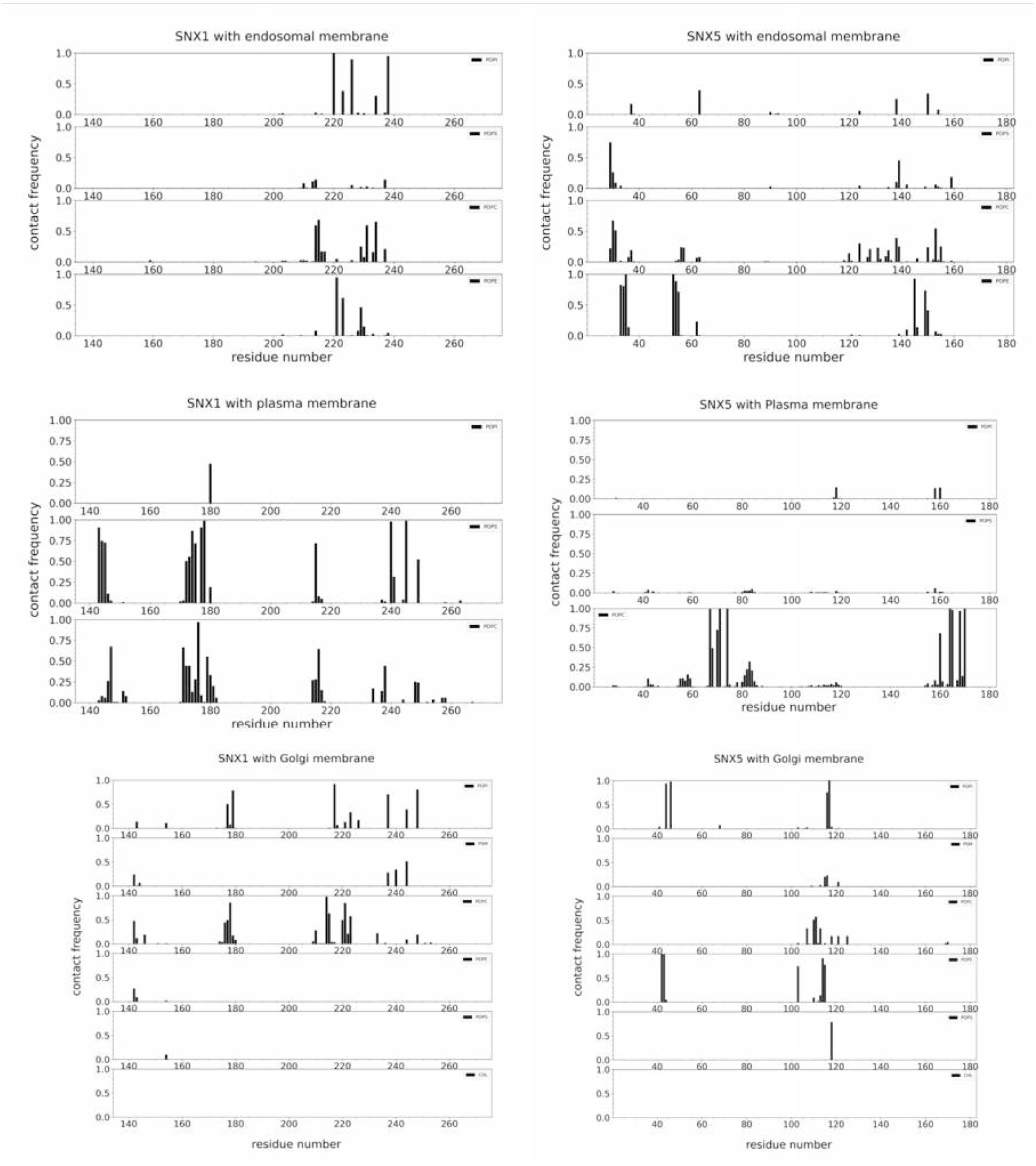
Residue-level contacts between SNX-PXDs and lipids in the three different membrane compositions. The left panel shows lipid interactions with SNX1, while the right panel shows interactions with SNX5. X-axis represents residue numbers, and y-axis indicates the contact frequencies.

In contrast to SNX1, the electrostatic potential of SNX5 clearly shows the lack of positively charged regions on the surface with an exception of the C-terminal end of *α*2 (Fig. S1 (B) in SI). A predominantly negatively charged region is present at the *α*1 and at the extended *α*2, which is the 15 residues insertion in SNX5-PXD that is absent in SNX1-PXD. Due to these differences in the surface electrostatic potential, the interaction of SNX5-PXD with the membrane would be different from SNX1. Hence, to understand its membrane binding property, we performed simulations of SNX5-PXD with the three membrane compositions similar to SNX1-PXDs. The Z-distance and contact analysis show that the SNX5-PXD inserts in the endosomal membrane, particularly the *β*1, *β*2-*β*3 loop and the loop in *α*2 double helix Fig.3(E). Notably, the hydrophobic residues VAL35, LEU55 and VAL155 show interaction with the POPE lipids (Fig.5). When simulated with the plasma membrane, the SNX5-PXD does not show any insertion Fig.3. However, the *α*1 and *α*2-*α*3 loop regions show interaction with POPC lipids involving the residues ARG67, TRP74, VAL165 and GLN170 (Fig. S5). Similarly, there is no insertion of PXDs in the Golgi membrane, while the loop in the double helix interacts with the lipids (Fig. S5). Particularly, the residues LYS44 and LYS117 interact with the PIP lipid and ARG42 interacts with the POPE lipid. These results show that SNX5-PXD expresses minimal interactions with the membrane irrespective of the composition. Membrane juxtaposed localization of SNX5-PXD is required for the function of the full SNX1-SNX5 heterodimer and these interactions are essential for SNX5-PXD to remain at the vicinity of the membrane. To further validate this study we have taken the membrane-bound orientations and performed REHT-PMP simulations, further simulation details are given in methods 0.2. We have taken last 50 ns of the all the replicas for further analysis. In these runs, the binding orientations of SNX1-PXD was similar as the classical MD simulations with endosomal and plasma membrane, which shows that these interactions are stable. We see a similar trend for SNX5-PXD runs as shown in Fig. S2 in SI. Together, our simulations clearly show that SNX1-PXD interacts strongly with the membrane when compared to SNX5-PXD. In order to further understand SNX1 behaviour, we designed mutants of SNX1-PXD and studied its interaction with the endosomal membrane.

Since the surface of SNX1-PXD is distributed with positively charged residues, and we show multiple binding orientations are possible, we performed mutagenesis studies to explore the most preferred binding orientation. We observed from our simulations that LYS223 interacts consistently with the three different compositions, and hence, we first modeled the LYS223ALA system with the endosomal membrane. Fig. S3 shows that the residues in the PPK loop continue to interact with the membrane; however, the PXD does not insert into the membrane (Fig.3(F)). Next, we included LYS214 to prevent the PPK loop interaction. Interestingly, in this system, the *β*2-*β*3 loop interacts with the membrane albeit without any insertion (Fig. S3 and Fig.3(F)). Among them, the residues 173-181 show strong interactions with the PIP and POPC lipids, and hence, we chose ARG177 and LYS179 for further mutation. Fig. S3 shows that mutating either ARG177ALA or LYS179ALA led to the recovery of PPK loop interactions. Interestingly, LYS179ALA mutation retained the interaction by the neighboring ARG177, while the vice versa was absent indicating the importance of ARG177. Hence, we modeled an SNX1-PXD with the endosomal membrane combining all these mutations, LYS223ALA, LYS214ALA, LYS221ALA and ARG177ALA. Surprisingly, even though the dominant positively charged regions were all mutated, the other sparsely distributed LYS and ARG residues in the *β*1 and *β*2 sheets were able to interact with the membrane in a limited manner without insertion. This further validates our assumption that the SNX1-PXD can adopt multiple orientations on the membrane according to the membrane composition.

The isolated PXDs might interact differently with a membrane when they are modeled independently. In the presence of the BAR domain, their dynamics might be restricted. Hence, to explore their binding in a full-length state, we modeled a SNX1-SNX5 heterodimer (discussed in the Methods section 0.1) on an endosomal membrane and performed all-atom simulations for a period of 2 microseconds (see movie file movieS2.mpeg in SI). It is assumed that the BAR domain of SNX-BAR proteins can assemble into higher order oligomers and induce membrane curvature. Our simulation has shown that a single heterodimeric unit of SNX1-SNX5 was unable to induce membrane curvature due to unstable oligomer interfaces. However, Z-distance analysis shows that the amphipathic helices of PXDs insert into the membrane while the BAR domain remains stable with peripheral interaction (Fig.4). Though the SNX1-PXD does not show deeper insertion into the membrane, its orientation is similar to the conformation seen in a cryo-EM study. Recently, the supra-molecular arrangement of SNX1-SNX5 heterodimers on a tubule of endosomal membrane composition was identified by cryo-EM. Though the cryo-EM maps were of a low resolution (~ 10 Å), this study has provided several insights into the membrane binding and remodeling ability of SNX proteins. They have clearly shown the role of heterodimer formation in membrane remodeling and cargo recognition. They have shown the favorable arrangement of PXDs at the sides of BAR domains and they are also involved in higher order assemblies. However, due to the low resolution of the cryo-EM densities, it is possible for the PXDs to adopt different orientations and yet fit in the cryo-EM densities. Through our simulation studies, we observe the binding of several residues in different membrane binding orientations and hence, we tested the possibility of fitting these orientations in the available cryo-EM map. Fig.4(G-H) shows that more than one orientation of the PXD is possible and this flexibility arises due to the presence of a flexible interdomain linker which also contains the amphipathic helix.

### Aromatic rich *π-π* interactions stabilize the SNX5-PXD and CI-MPR complex

The crystal structure of SNX5-PXD in complex with CI-MPR tail region has provided two different binding modes and we wanted to model the dynamics of the complex to arrive at the stabilizing interactions. Hence, we performed a well-tempered metadynamics simulation of SNX5-PXD with the CI-MPR tail to extract the binding free energy profile of the complex where we explored several binding poses including the one available in the crystal structure (PDB ID: 6N5X). In addition, we modeled a complex of SNX1-PXD with CI-MPR as well to understand why SNX5-PXD preferentially bind to CI-MPR as compared to SNX1-PXD. Initially, we identified the binding pockets in SNX5-PXD and SNX1-PXD by performing binding pocket analysis using CASTpFold (54). The major binding pocket in SNX5-PXD is at the site of CI-MPR binding in the crystal structure, which includes the site between the *β*1-*β*2 loop and the *α*2 double helix. In contrast, the largest binding pocket in SNX1-PXD is considerably smaller than SNX5-PXD, and buried between the helices and the PPK loop, which is not equivalent to the CI-MPR binding site in SNX5-PXD (Fig.6 (A-B)). The collective variable was defined as the distance between LEU2370 of CI-MPR and PHE136 of SNX5, and the structurally equivalent LEU235 in the SNX1-PXD. Metadynamics simulations were performed until convergence is achieved (see SI Fig. S4 and Methods for detailed information 0.4), and the converged free energy surface for CI-MPR binding with SNX1-PXD and SNX5-PXDs are shown in Fig.6. All conformations comprising the minimum energy basin were extracted and aligned in Fig.6 to depict the binding modes sampled by CI-MPR during the metadynamics simulation.

**Figure 6:**
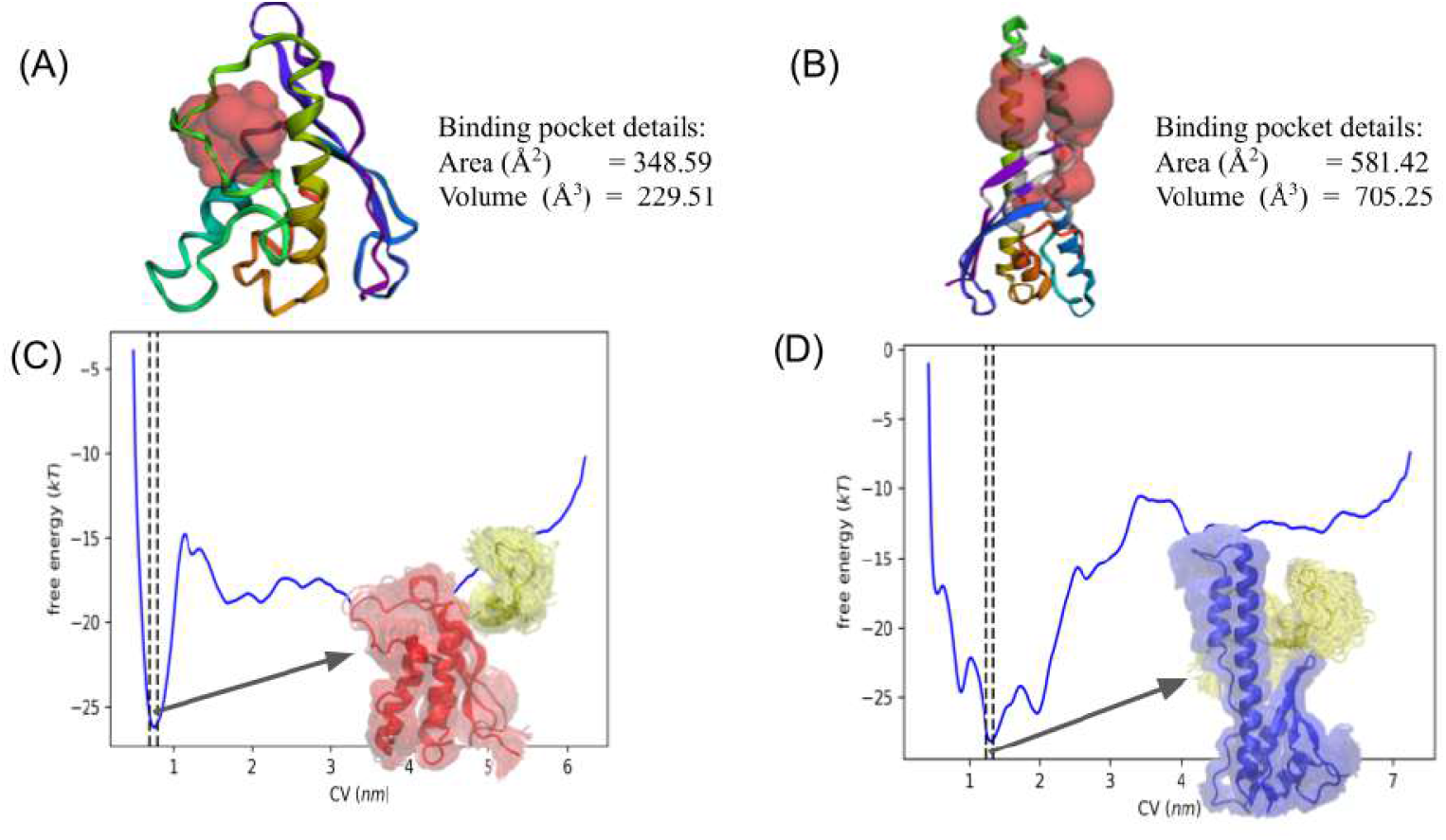
(A, B) SNX1-PXD and SNX5-PXD binding pocket analysis done using CASTpFold, which shows larger binding pocket in SNX5-PXD compared to SNX1. (C, D) shows the free energy landscape of SNX1-PXD and SNX5-PXD with CI-MPR tail region. X-axis shows the CV distance and y-axis shows the free energy. Conformations that contribute to minimum energy are aligned and shown in cartoon representation for both SNX1-PXD and SNX5-PXD systems, where SNX1-PXD, SNX5-PXD and CI-MPR tail are shown in Red, Blue and Yellow, respectively.

Though dissociation free energies for CI-MPR binding with SNX1 and SNX5 were highly similar, the potential of mean force (PMF) shows an intermediate high energy barrier for association with SNX1-PXD when compared to the favorable monotonic binding pathway in SNX5-PXD. The CI-MPR association with SNX5-PXD is more kinetically accessible than that with SNX1-PXD. Contact analysis of these complex clusters showed a larger number of electrostatic interactions with SNX5-PXD (Fig.7 (A)) and a comparatively lesser number of interactions with SNX1-PXD (Fig. S5). The interacting pairs of CI-MPR with SNX5-PXD present in the crystal structure were dynamic during simulation due to the presence of other aromatic and charged residues at the vicinity (Fig.7 (A)). Particularly, the stacking interaction between PHE136 and CI-MPR (LEU2370/MET2371) seen in the crystal structure was replaced by stacking interaction with TYR132 and TYR2353 or TRP2369. In addition to this, several *π*-stacking interactions and salt bridges were observed between SNX5-PXD and CI-MPR, indicating the dynamic nature of their binding (Fig.7 (B-E)). Particularly, the CI-MPR residues TYR2351, TYR2353 and TRP2369 involve in *π*-*π* stacking and T-shaped stacking with PHE99, TYR132 and PHE136 of SNX5-PXD (Fig.7). In contrast to these strong electrostatic interactions in the SNX5-PXD, the SNX1-PXD is dominated by non-specific and transient hydrophobic interactions involving residues like LEU235, PRO152, VAL163 and GLU236 among others (Fig. S5 in SI). These results clearly indicate that SNX5-PXD has the ability to adapt and strongly bind to CI-MPR (and possibly different cargo proteins with similar beta strand motif) due to the presence of aromatic and charged residues at the binding site.

**Figure 7:**
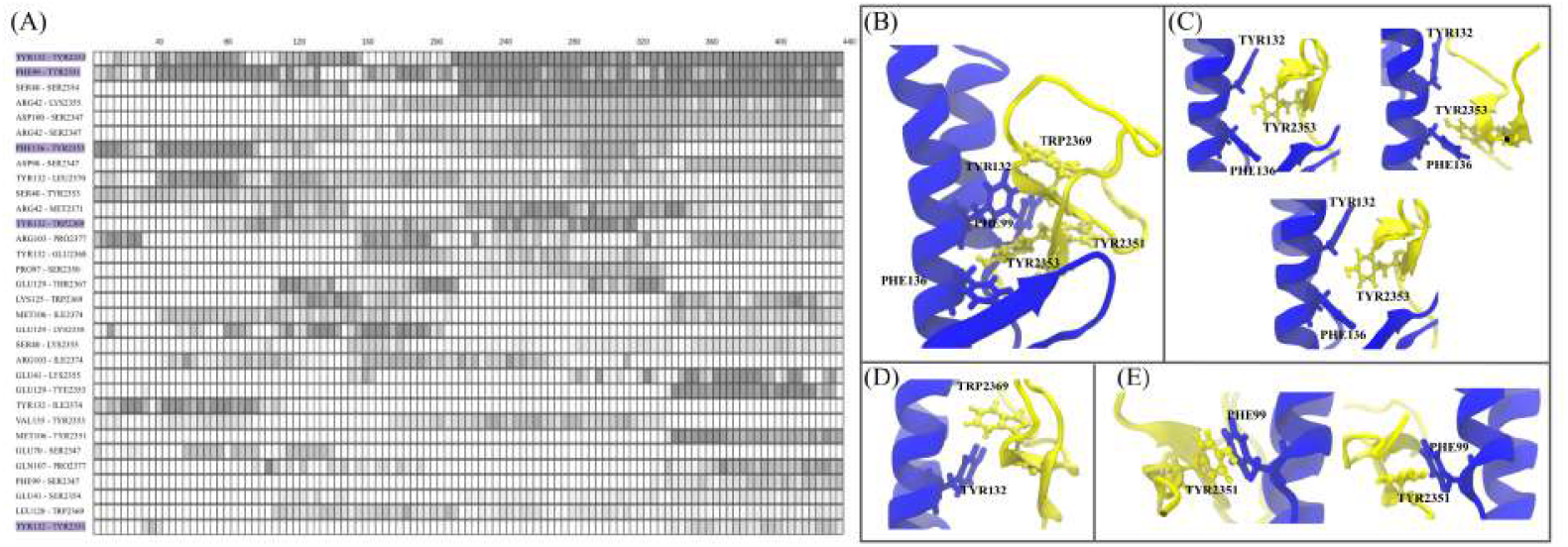
Contact analysis between SNX5-PXD and CI-MPR, extracted from the structures contributed to the minimum energy. (A) shows the lifetime of these interactions, with individual conformations on the x-axis (every 4th conformation is shown) and residue pair on the y-axis. (B) shows the important *π*-*π* interactions stabilizing these complexes. TYR2353 shows stacking and T-shaped interactions with TYR132 and PHE136, as depicted in (C). T shaped *π*-*π* interactions between TYR132 and TRP2369 is shown in (D) and Stacking and T shaped interaction between PHE99 with TYR2351 in (E)

### SNX1-SNX5 heterodimer optimizes endosomal tubulation and cargo sequestration

The mesoscopic continuum model allows us to feed in the interaction propensities between different components of the system and look at the emergent behavior at larger scales as a result of the interactions arising at molecular scales. As a way to highlight the scope of this method, we first present the effects on membrane remodeling and cargo sequestration under different combinations of interactions between SNX1, SNX5 and CI-MPR proteins (see Fig.8(A-C)). To reiterate, the Red nematics represent SNX1, Blue represents SNX5 and Yellow is for CI-MPR cargo protein. In Fig.8(A), we simulate the Hamiltonian under the condition that none of the proteins are interacting with each other. Based on the inputs from our molecular simulations, we chose to keep the parameters of continuum Hamiltonian such that SNX1 binds strongly to the membrane, SNX5 weakly interacts with the membrane and CI-MPR has no localization propensity vis a vis any other protein. SNX1 nematics form tubules and the tubules do not have SNX5 or CI-MPR proteins in them (Fig.8(A)). When we change the conditions that allows interactions between SNX1 and SNX5 (a theoretical construct that models the possibility of heterodimer formation), we again see formation of tubules and ridges that are rich in SNX1-SNX5 but no CI-MPR sequesteres in the remodeled tubules. Since we did not input any preference for CI-MPR to bind to the curvature generating in-plane proteins, the CI-MPR nematics (yellow) energetically prefer the low curvature non-tubular morphologies. In Fig.8(C), we change the parameters such that CI-MPR has affinity for SNX5 and it can be clearly seen that we do have both tubulations as well as cargo sorting within these tubes (see movie file movieS3.mpeg in SI). Interestingly, though at face value, it seems that CI-MPR may not have any effect on membrane remodeling, we observe that at higher densities and varying degree of interaction affinity with SNX5, CI-MPR does affect the tabulation patterns (Fig. S6). When we simulated our system with densities or surface coverage of SNX1 and SNX5 at 25% each while changing the CIMPR densities to three different values (5%, 10% and 20%), we see differences in tubule formation patterns (see Fig. S6 (A-C)), which points to the non-linear effect of cargo incorporation in the tube on membrane tubulation propensities.

**Figure 8:**
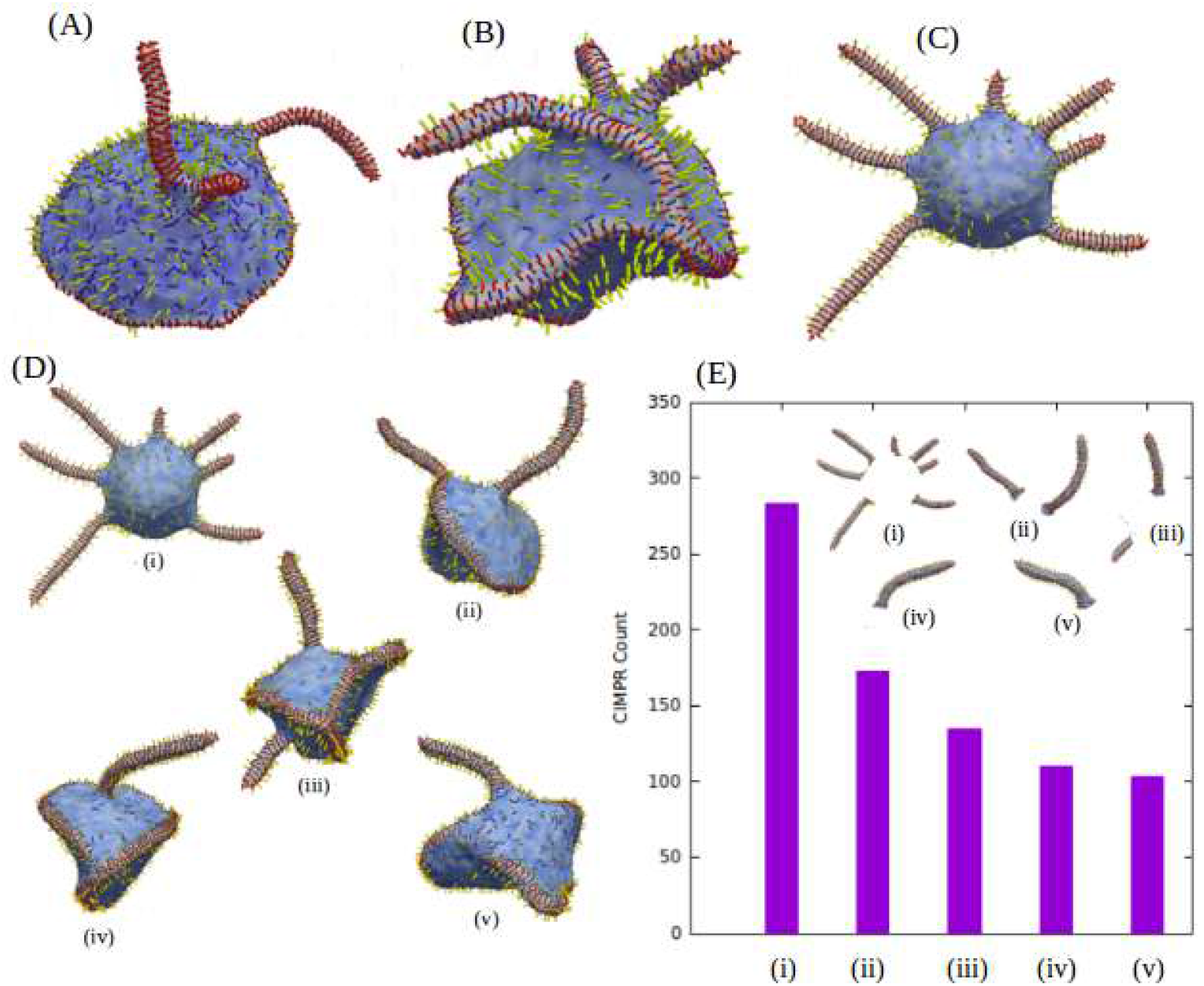
Effect of interplay with the interaction between SNX1, SNX5, and CI-MPR are tested, possible interactions defined are the possibility of SNX1 homodimerization, SNX1-SNX5 heterodimerization and SNX5-CI-MPR which helps to transport cargo. In (A) we allowed only SNX1 homodimerization, (B) shows membrane remodeling when hetrodimerization of SNX1-SNX5 is allowed and CI-MPR binding to SNX5 is zero. (C) shows how membrane remodeling and cargo transportation happens when hetrodimerization and cargo recognition is allowed. (D) Variation in the deformed shapes due to the local densities of SNX1 and SNX5. Total density of SNX1 and SNX5 is 50% but the membrane shapes are changing due to local variation in the densities of SNX1 and SNX5. Here, the densities of SNX1 and SNX5 are (35%, 15%),(30%, 20%), (25%, 25%), (20%, 30%) and (15%, 35%), respectively in (i)-(v). (E) Histogram shows the CI-MPR count in the tube region which is transported by the SNX1-SNX5 hetrodimers. Other parameters are *ϰ*= 20, *ϰ*_∥_ = 50 (SNX1) and 5 (SNX5) and *ϵ*_*LL*_ = 3 (SNX1-SNX5) and 1.5 (SNX1-SNX1) in *K*_*B*_*T* unit and the value of *C*_∥_ is 1.0 for both SNX1 and SNX5.

We also explored the tubulation versus cargo sorting propensities with simulating use cases where different levels of cross-interactions were tried between SNX1 and SNX5. The systems with higher interactions between SNX1 and SNX5 led to formation of tubes that had CI-MPR cargo sorted into the tube (Fig. S7(A, B)). Since SNX5 is known to be not the primary remodeling protein, we also tried simulations with high SNX1-SNX5 interactions but different levels of SNX5 binding (and curvature-inducing ability) to the membrane. We can see that the propensity to form tubes reduces with a much weaker SNX5 membrane association though cargo sequestration to the tube did not get affected as much. This difference is shown in Fig. S7(A) and Fig. S7(B). Also, when we changed the SNX1 and SNX5 cross interactions such that SNX1 homodimer formation tendencies outweighs the SNX1-SNX5 heterodimer formation (Fig. S7(C) and Fig. S7(D), respectively), we observe that the tubes are formed due to SNX1 homodimer and cargo sorting of CI-MPR into the newly formed tubes is largely compromised. It is interesting to see that though the curvature of SNX1 and SNX5 are thought to be similar, SNX1-SNX5 heterodimer has a weaker membrane bending ability than SNX1 homodimers. This is reasoned from the point of view of higher membrane affinity of SNX1, which provides the energy to overcome the resistance to bending for the membrane. In order to test this comprehensively, we studied the effect on membrane tubulation and cargo sorting due to different membrane binding and curvature propensity. In Fig. S8, we show membrane deformation results with the different values of membrane-protein interaction and radius of curvature of SNX1 and SNX5. The ratio of induced membrane-protein rigidity of SNX1 and SNX5 varies on the Y axis and the radius of curvature of SNX1 and SNX5 varies on the X axis. Deformation of the membrane increases when both SNX1 and SNX5 monomers interact with the membrane and the curvatures of both proteins are the same. Besides the cross-interactions, the relative populations of SNX1 and SNX5 plays a major role in driving the balance between tubulation and cargo sorting. We ran simulations where we kept total combined density of SNX1 and SNX5 to be 50% and looked at the counts of CI-MPR sorting into the tubes. Interestingly, increasing SNX5 does not always lead to higher number of cargoes getting sorted in the tubes since higher SNX5 densities compromises tube formation. In Fig.8 (D-E), we report data from five runs with following densities of SNX1 and SNX5: (i) SNX1 35% and SNX5 15%, (ii) SNX1 30% and SNX5 20%, (iii) SNX1 25% and SNX5 25%, (iv) SNX1 20% and SNX5 30%, and (v) SNX1 15% and SNX5 55%. The total CI-MPR counts in the tubes show a steady decline (Fig.8 (E)) despite no changes in interaction between SNX5 and CI-MPR and an abundance of CI-MPR on the membrane surface away from the tubes. This is due to less tubes getting formed when SNX1 is reduced, which is the primary membrane remodeling component. It is clear that with incorporation of SNX5 as a heterodimer, the heterodimer is able to sort cargoes but the tubulation ability of SNX1 is mitigated to some extent. Together, our mesoscopic simulations show that the SNX1-SNX5 heterodimer has evolved such that both tubulation and cargo-sorting can take place albeit with a constraint on the extent of tubulation as well as cargo sorting in this retromer-independent trafficking pipeline.

## CONCLUSION

Sorting Nexins are a family of proteins responsible for endosomal sorting and transport of cargo proteins, recycling them between Endosomes, Plasma membrane and Golgi bodies through the Trans-Golgi Network. In concert with the larger retromer complex, these proteins mainly bind to membrane, accumulate cargo and remodel the membrane to form tubules or vesicles for transport. However, recent discoveries, especially in the ESCPE-1 pathways, point to distinct functional roles for SNX-BAR dimers and retromers (2, 19). In this study, we have focused on the retromer-independent SNX1-SNX5 heterodimeric systems and explored the molecular and thermodynamic driving forces leading to specific membrane and cargo molecular recognition as well as larger scale cargo sorting and endosomal tube formation.

Specific phosphoinositols (PIPx) lipid recognition by various PXD has been a subject of intense discussion in recent literature (8, 12, 13, 20, 55). Since a specific PIP lipid type can be found on different organelle membrane, there seems to be a paradigm shift in the understanding where single PIP lipid is now not reliably considered as a specific recognition motif or organelle membrane marker for important membrane adaptor domains such as PXDs, PH Domains, FYVE and other domains. It is possible that a distinct set of lipids around the given PIP lipid forms an oligomeric functional nanodomain that acts as a “molecular portrait” (56) or “lipid fingerprints” (57) that are collectively recognized by the membrane adaptors (55, 58–62). Our all-atom simulation data, where we modeled the membrane binding of isolated SNX1-PXD and SNX5-PXD with three different membrane composition, reveals different membrane binding behavior. Until recently, the membrane binding property of SNX1 PX domain, which is the primary membrane recognition adaptor in the heterodimeric SNX1-SNX5 complex, was considered to be restricted to two primary binding sites. In our work, we clearly see that the SNX1 PX domain can explore several additional lipid binding sites, among which the residues LYS223, LYS214, LYS221 and ARG177 are important. Moreover, we see that different segments of the protein has preferred association with different membrane, which suggest the different ways in which specific organelle recognition may be taking place where structural motifs on adaptor and local lipid nanodomain organization cooperate for binding.

Detailed all-atom simulations and Metadynamics based free energy calculations also shed some important insights into the molecular determinants behind the specific cargo recognition ability of SNX5-PXD. Contrary to the enriched positively charged residues in SNX1, the SNX5 PX domain contains several hydrophobic and aromatic residues, which form a chemical motif for recognizing cargo. This includes the aromatic *π*-*π* and T-shaped stacking of PHE99, TYR132 and PHE136 with the CI-MPR residues as shown in our study. From this study, we clearly showed how SNX1-PXD and SNX5-PXD behave differently with endosomal membrane, plasma membrane, Golgi membrane and CI-MPR.

Along with atomic-level mechanistic insights, we also modeled the transport of CI-MPR by SNX1-SNX5 heterodimers using mesoscopic modeling. Cryo-EM studies have shown the oligomerized state of SNX1 homodimers and SNX1-SNX5 heterodimers collars over a tubule (14,22). Our simulations show the mechanisms of cargo sorting and transport into tubules by modeling organelle-level systems using the mesoscopic dynamically triangulated surface model. Here, we used several hypothetical use cases mimicking various mechanistic possibilities to thermodynamically establish the pathway to cargo sorting and tubule formation. Our mesoscopic data clearly shows that the heterodimeric SNX1-SNX5 causes concomitant cargo sequestration and tubulation in a retromer-independent manner. This is interesting since we could show that SNX1 homodimers could form tubes but not sequester cargo, while use cases where SNX5 in SNX1-SNX5 heterodimer was not allowed to interact with the CI-MPR proteins, tubulation and cargo sorting in tubes were enormously compromised. Our study provides important insights into the membrane remodeling by SNX proteins and the need for heterodimerization, which plays a key role in cargo transport when the process has to take place without the retromer complex.

## DATA AVAILABILITY STATEMENT

Our codes, models, all the inputs files and trajectories of all the simulations are publicly available at the Zenodo portal, MultiscaleSims-SortingNexins.

## AUTHOR CONTRIBUTIONS

AS conceived and designed the research. SCD and GK performed the research. Data analyses was carried out by all authors. Continuum mechanics modeling was carried out by GK while SCD generated data related to molecular simulation and also contributed to Continuum mechanics simulations. SCD and GK wrote the first draft of manuscript. AS polished the draft with help of all authors.

## ACKNOWLEDGMENTS

The authors thank Sangeetha Balasubramanian for critically reading the manuscript and suggesting changes. SCD thanks the Ministry of Education for his graduate fellowship, and GK thanks the Axis Bank Postdoctoral Fellowship at the Indian Institute of Science (IISc), Bangalore for financial support. Financial support from the IISc and the high-performance computing facility “Beagle” setup from grants by a partnership between the Department of Biotechnology of India and the Indian Institute of Science (IISc-DBT partnership program) is greatly acknowledged. AS thanks DST for the National Supercomputing Mission grants (DST/NSM/R&D-HPC-Applications/2021/03.10, DST/NSM/R&D-HPC-Applications/Extension Grant/2023/27). AS also acknowledges the FIST program sponsored by the Department of Science and Technology, India, which supports the MBU infrastructure. AS would also like to thank the Teams Science Grant from the DBT-Wellcome Trust India Alliance (Grant number: IA/TSG/21/1/600245). AS also thanks the DBT National Network Project (NNP) grant (BT/PR40323/BTIS/137/78/2023) and the Matrics grants (MTR/2023/001040) from the Science and Engineering Board (SERB), India.

## SUPPLEMENTARY MATERIAL

Document S1

Supplementary information contains eight figures (Fig. S1 to Fig. S8) and three movie files.

Movie file 1 (movieS1.mpeg)

Fitting of SNX1-SNX5 dimer modelled using membrane bound SNX1-PX domain orientation shown in red. For better visualization other domains of this structure is not shown. Structure of full lenghth SNX1-SNX5 heterodimer (pdb id 8AFZ) is shown in tan and cryo-EM map is shown in grey.

Movie file 2 (movieS2.mpeg)

SNX1-SNX5 full length protein on lipid bilayer with endosomal membrane composition. SNX1-PXD, SNX5-PXD, amphipathic helix and BAR domains are shown in Red, Blue, Magenta, and Green colors, respectively. For endosomal membrane, the lipids PC, PE, PS, PI3P, and PI(3,5)P are represented as Grey, Orange, Green, Magenta, and Red, respectively

Movie file 3 (movieS3.mpeg)

This shows the membrane remodeling due to SNX1-SNX5 heterodimerization and sorting of CI-MPR to the endosomal tubes using mesoscale modeling simulation. In this SNX1, SNX5 and CI-MPR are shown in Red, Blue and Yellow, respectively.

## Supporting Information

**Fig. S1:**
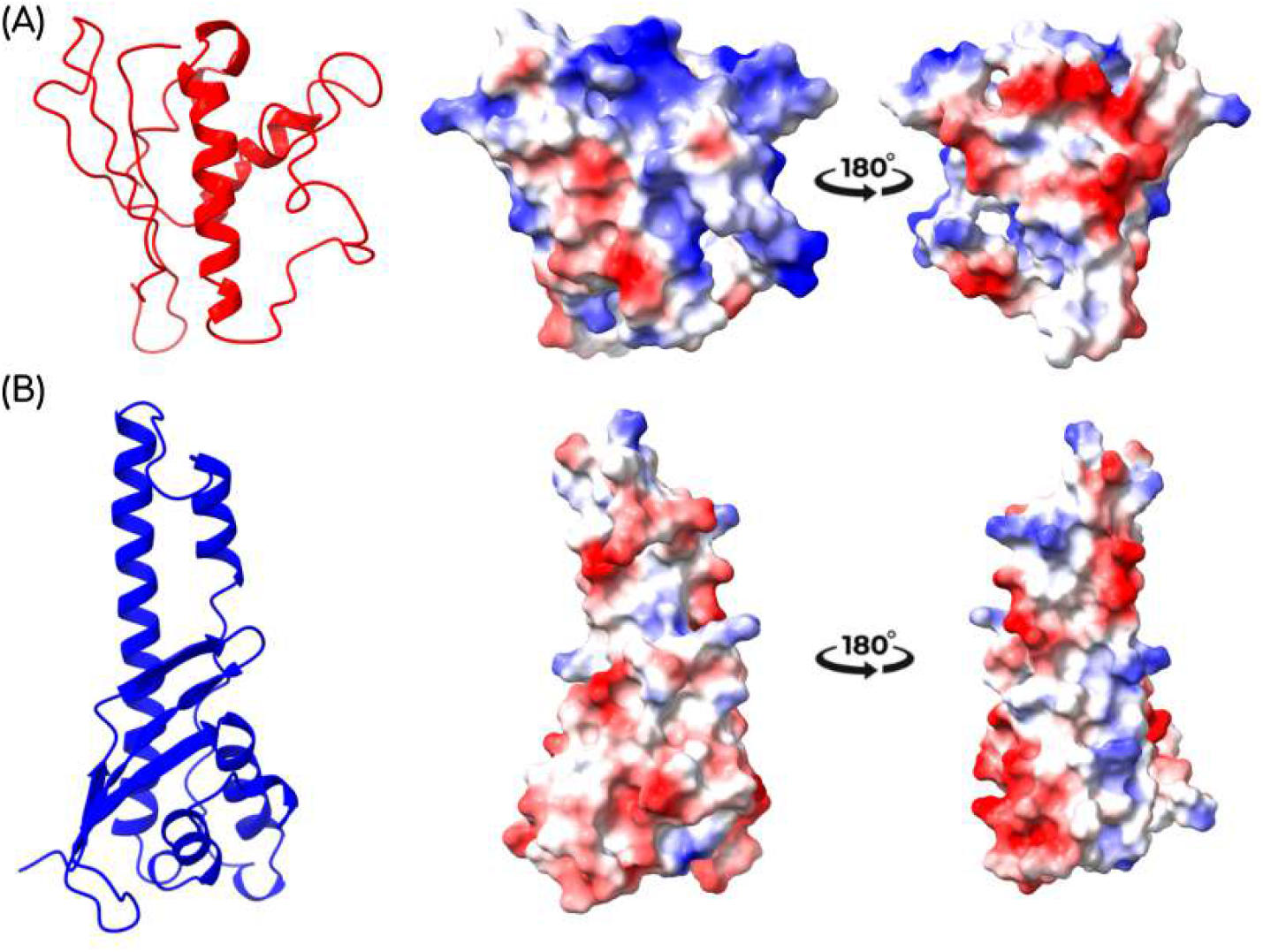
Electrostatic potentials of SNX1-PXD (A) and SNX5-PXD (B) depicting the distribution of positive and negatively charged residues on their surface.

**Fig. S2:**
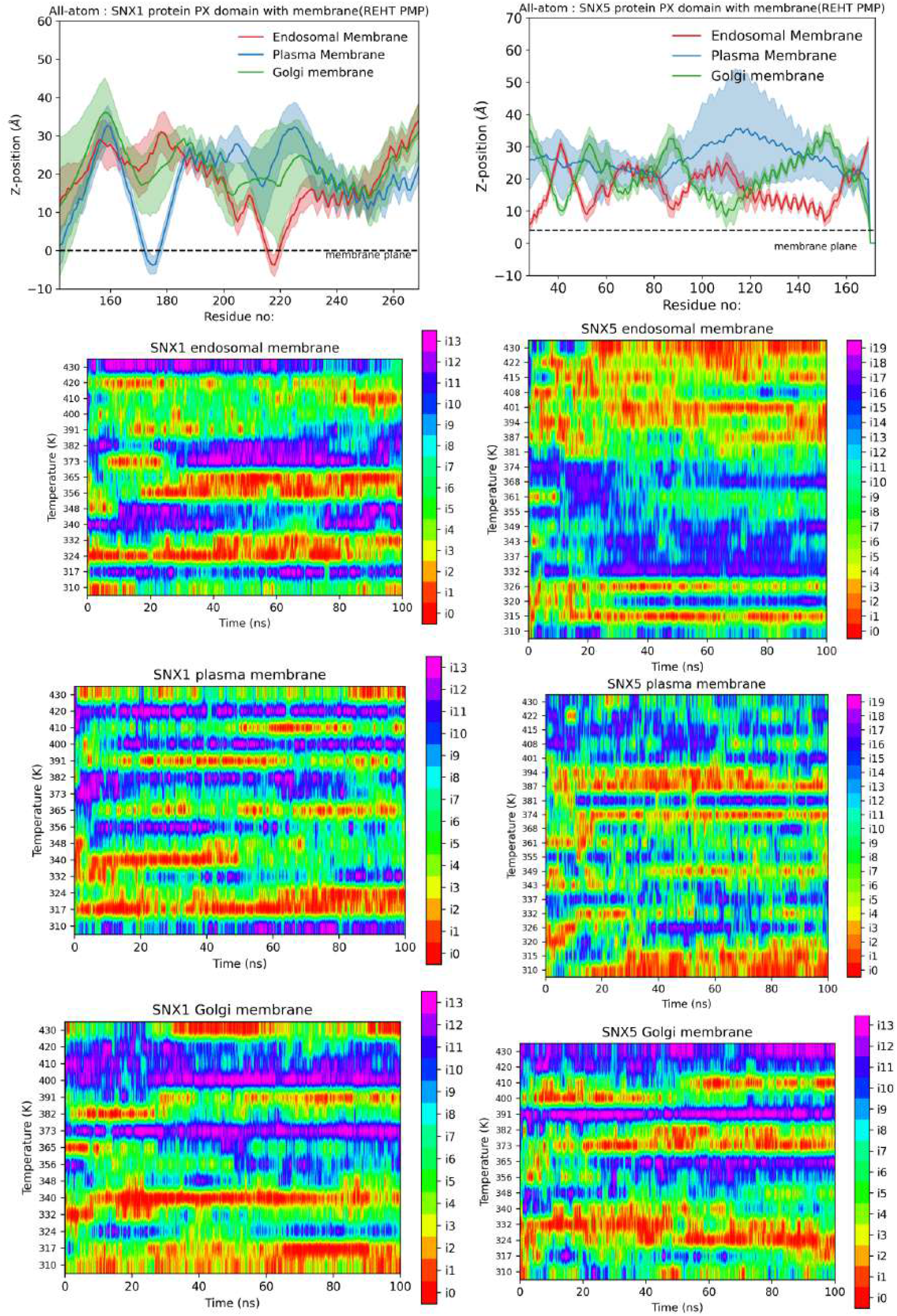
The top panel shows the z-distance between the protein and the membrane, with SNX1 shown on the left and SNX5 on the right. The remaining panels illustrate the exchange patterns extracted from the REHT-PMP replicas.

**Fig. S3:**
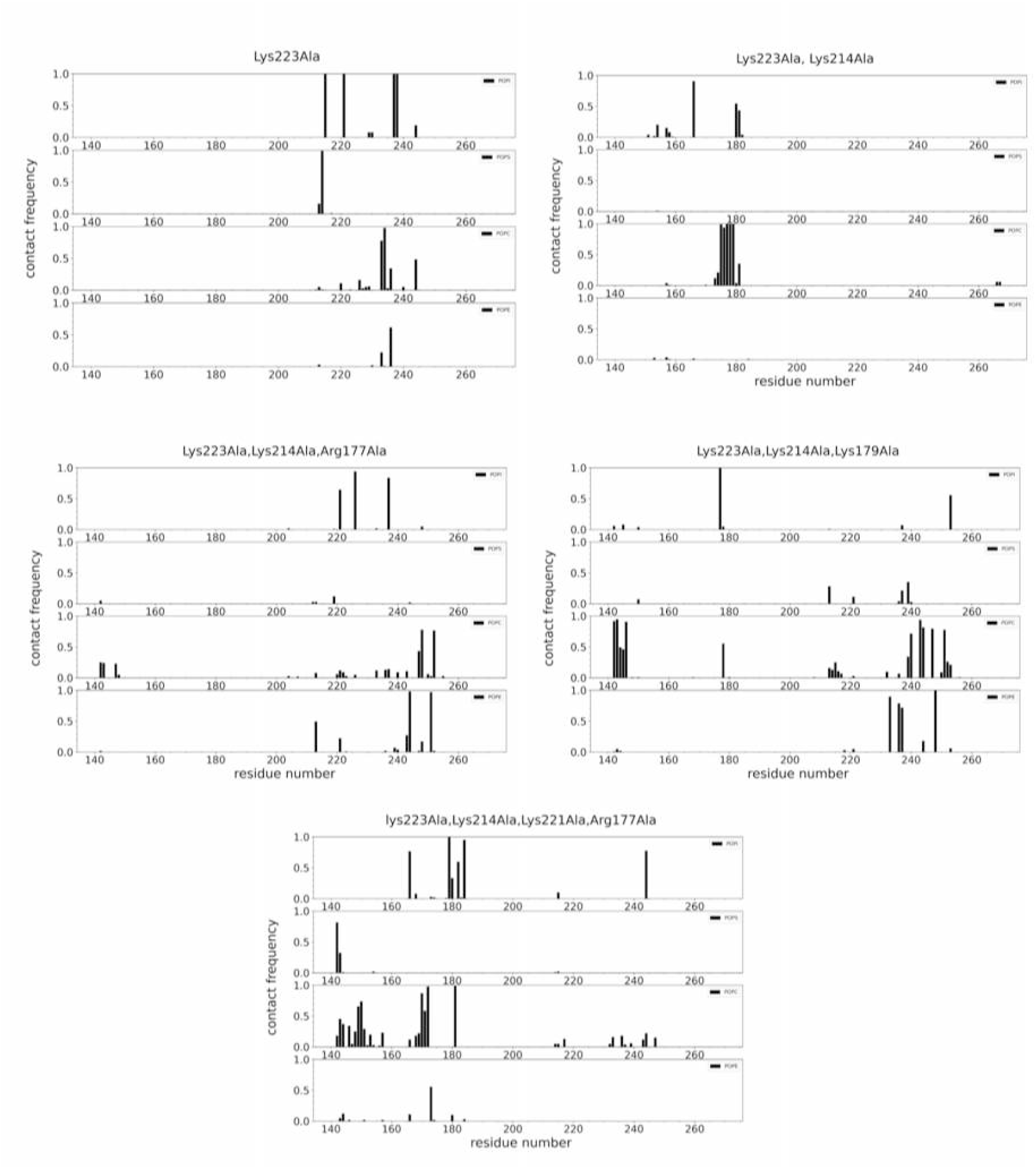
Residue-level contacts between SNX1-PXD and endosomal membrane with different mutations. X-axis represents residue numbers and y-axis indicates the contact frequencies.

**Fig. S4:**
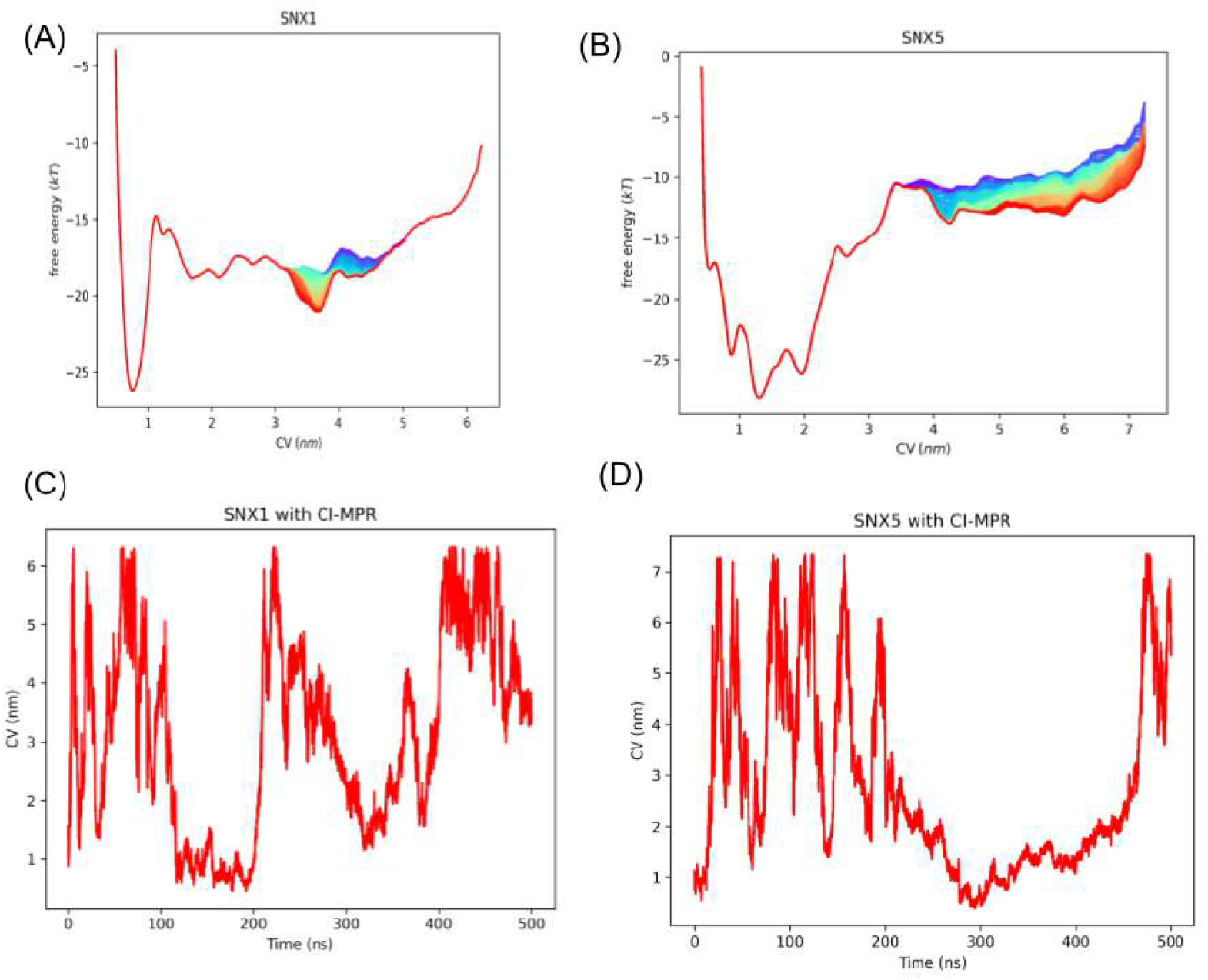
Convergence analysis of well-tempered metadynamics runs. The free-energy profiles and the time evolution of CVs for SNX1 (A, C) and SNX5 (B, D) is shown.

**Fig. S5:**
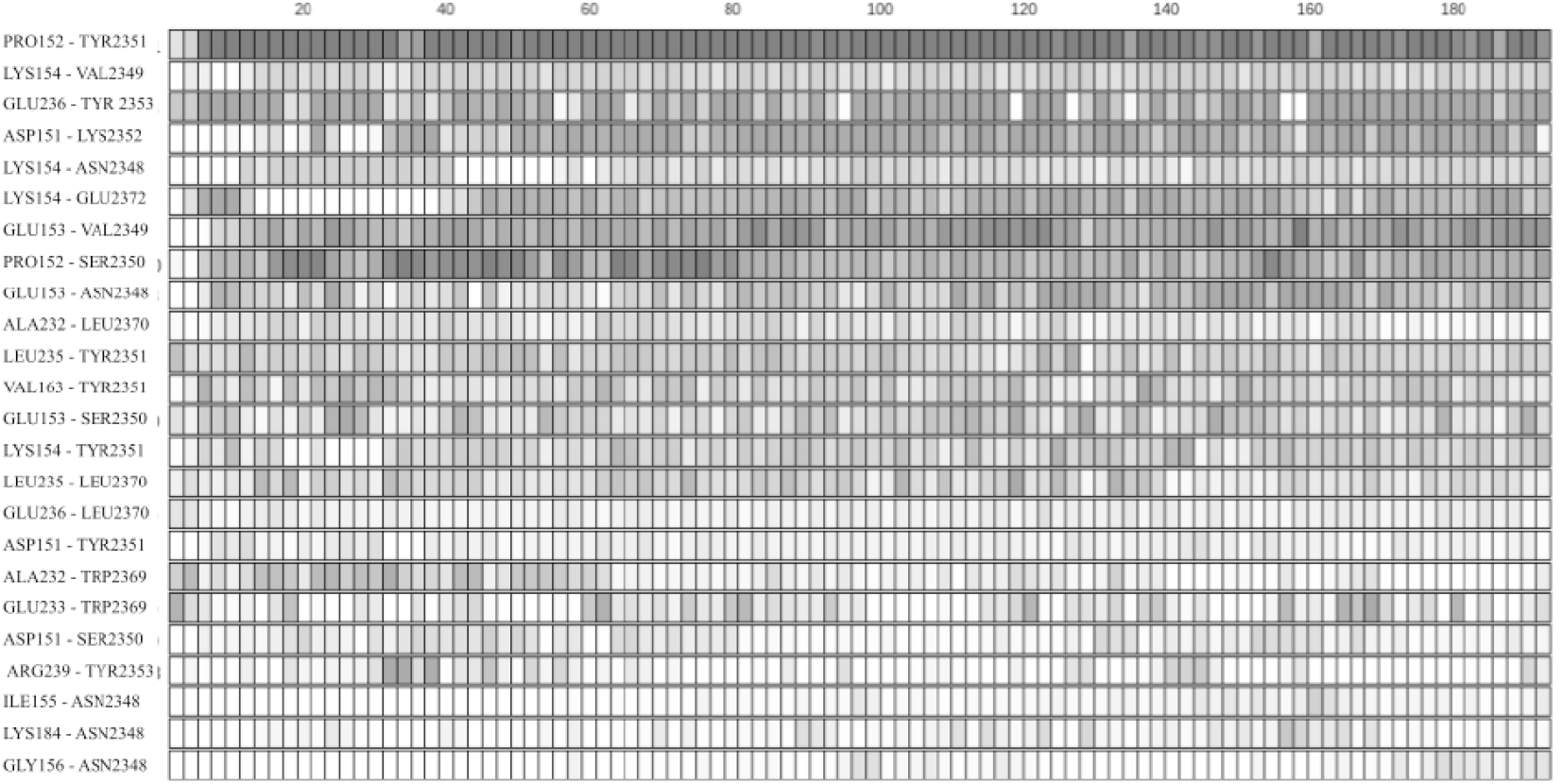
Contact analysis between SNX1-PXD and CI-MPR conformations which contribute to the minimum energy. This shows the lifetime of these interactions, with individual conformations on x-axis (every 4th conformation is shown) and residue pairs on y-axis.

**Fig. S6:**
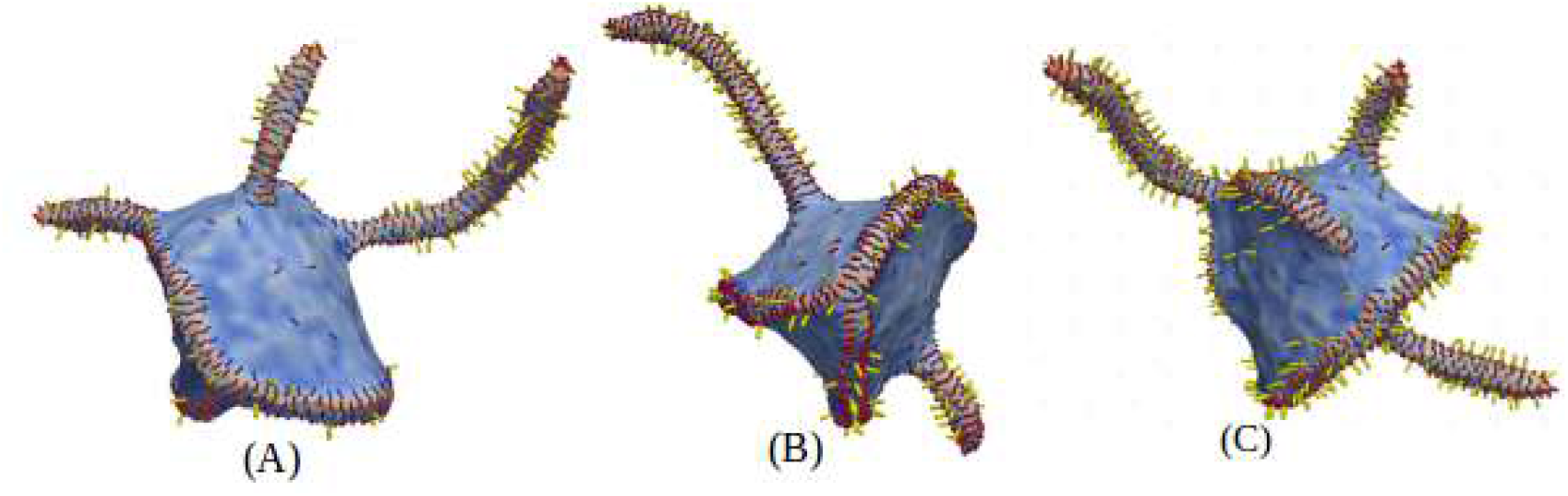
Different membrane morphologies due to variation in the density of CI-MPR. Here the densities of SNX1 and SN5 monomers are 25% and the densities of CI-MPR in A-C are 5%, 10% and 20% respectively. Other parameters are *κ* = 20, *κ*_∥_ = 50(SNX1) and 5 (SNX5) and *ϵ*_*LL*_ = 3 (SNX1-SNX5), 1.5 (SNX1-SNX1) in *K*_*B*_*T* units and the value of *C*_∥_ = 1.0.

**Fig. S7:**
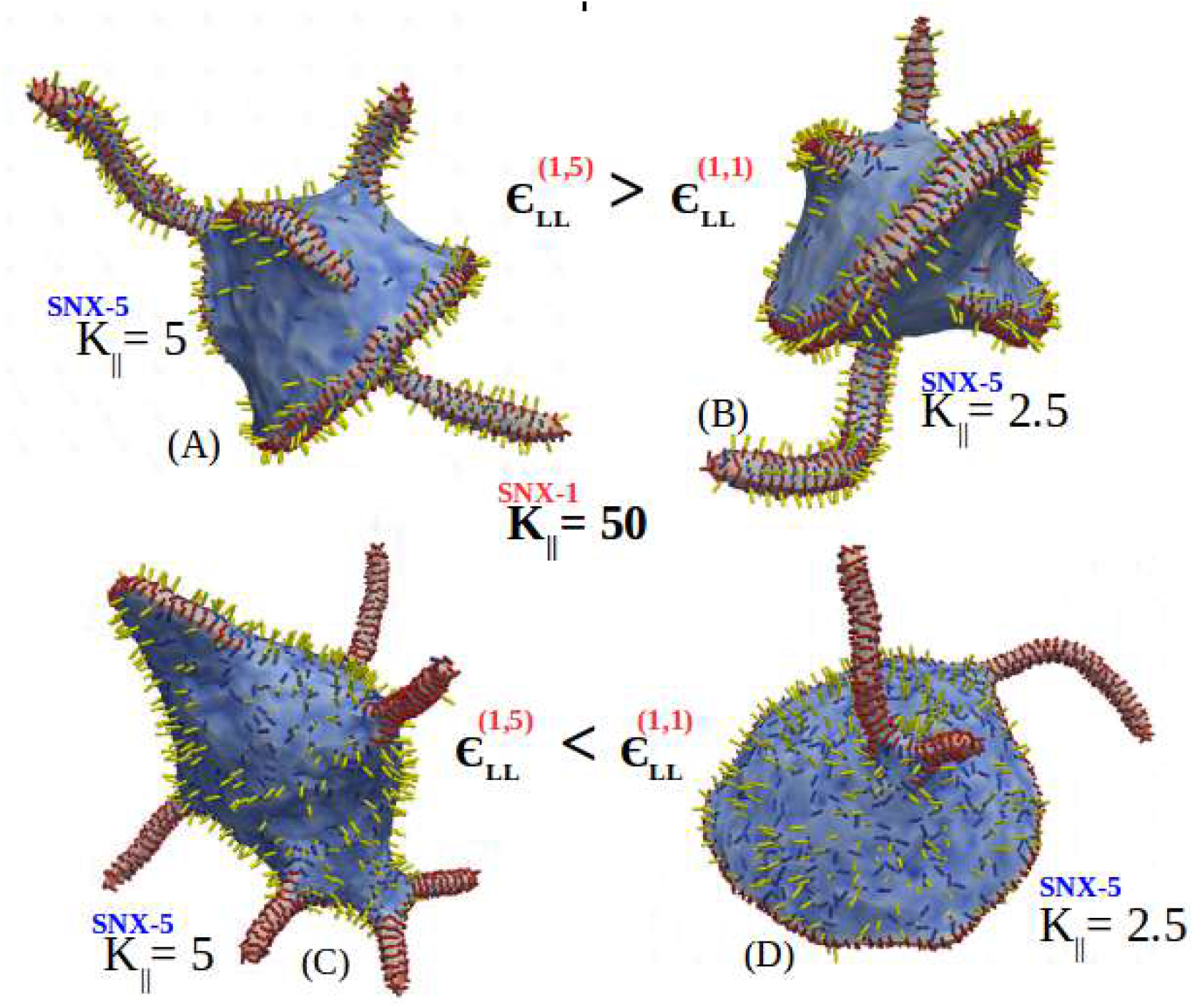
Membrane remodeling due to hetrodimers and homodimers of SNX1 and SNX5. In (A,B), heterodimers are formed due to high value of interaction between SNX1-SNX5 and shape is changing due to different values of induced bending rigidity due to SNX5 which is very low as compared to SNX1. Here maximum number of CI-MPR are transported in the tube. In (C,D), membrane remodeling due to SNX1 homodimers. In this case there is no transportation of CI-MPR in the tube. Here tubes are formed by the SNX1 homodimers and all the CI-MPR and SNX5 available on the bare membrane. Here the membrane bending rigidity of the membrane is *κ* = 20 *K*_*B*_*T, C*_∥_ = 1.0 and the densities of SNX1 and SNX5 are equal (25 %).

**Fig. S8:**
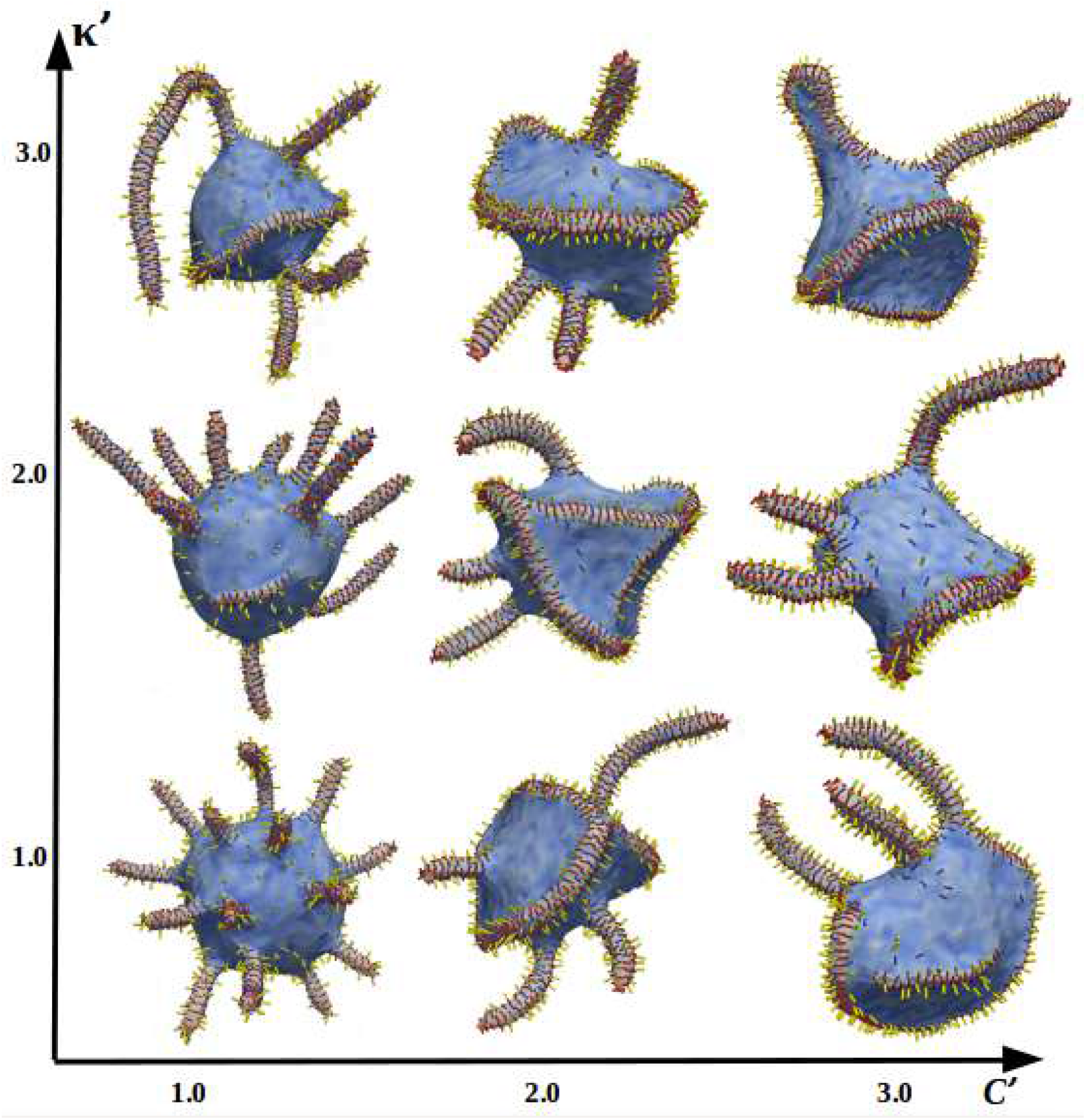
Membrane deformation results with the different values of the ratio of Induced membrane bending rigidities(*κ*_∥_) and radius of curvatures(*C*_∥_) of SNX1 and SNX5. The ratio of the radius of curvature(*C*^*′*^) of SNX1 and SNX5 varies on the x-axis and the ratio of induced membrane bending rigidity(*K*^*′*^) due to SNX1 and SNX5 varies on the y-axis. It shows the deformation of the membrane increases when both SNX1 and SNX5 monomers interact with the membrane and the curvature of both protein monomers is equal. Here the densities of SNX-1 and SNX-5 are equal(25 %). Other parameters are *κ* = 20, *ϵ*_*LL*_ = 3(SNX1-SNX5), 1.5 (SNX1-SNX1) in *K*_*B*_*T* units and the value of *C*_∥_ = 1.0.

## REFERENCES

1. Cullen, P. J., 2008. Endosomal sorting and signalling: an emerging role for sorting nexins. Nature reviews Molecular cell biology 9:574–582.

2. Simonetti, B., J. L. Daly, and P. J. Cullen, 2023. Out of the ESCPE room: Emerging roles of endosomal SNX-BARs in receptor transport and host–pathogen interaction. Traffic.

3. Weeratunga, S., B. Paul, and B. M. Collins, 2020. Recognising the signals for endosomal trafficking. Current opinion in cell biology 65:17–27.

4. Cullen, P. J., and F. Steinberg, 2018. To degrade or not to degrade: mechanisms and significance of endocytic recycling. Nature reviews Molecular cell biology 19:679–696.

5. Li, C., S. Z. A. Shah, D. Zhao, and L. Yang, 2016. Role of the retromer complex in neurodegenerative diseases. Frontiers in aging neuroscience 8:42.

6. Zhang, H., T. Huang, Y. Hong, W. Yang, X. Zhang, H. Luo, H. Xu, and X. Wang, 2018. The retromer complex and sorting nexins in neurodegenerative diseases. Frontiers in aging neuroscience 10:79.

7. Lauffer, B. E., C. Melero, P. Temkin, C. Lei, W. Hong, T. Kortemme, and M. von Zastrow, 2010. SNX27 mediates PDZ-directed sorting from endosomes to the plasma membrane. Journal of Cell Biology 190:565–574.

8. Chandra, M., and B. M. Collins, 2019. The phox homology (PX) domain. Protein Reviews–Purinergic Receptors: Volume 20 1–17.

9. Hanley, S. E., and K. F. Cooper, 2020. Sorting nexins in protein homeostasis. Cells 10:17.

10. Cheever, M. L., T. K. Sato, T. de Beer, T. G. Kutateladze, S. D. Emr, and M. Overduin, 2001. Phox domain interaction with PtdIns (3) P targets the Vam7 t-SNARE to vacuole membranes. Nature cell biology 3:613–618.

11. Xu, Y., H. Hortsman, L. Seet, S. H. Wong, and W. Hong, 2001. SNX3 regulates endosomal function through its PX-domain-mediated interaction with PtdIns (3) P. Nature cell biology 3:658–666.

12. Karathanassis, D., R. V. Stahelin, J. Bravo, O. Perisic, C. M. Pacold, W. Cho, and R. L. Williams, 2002. Binding of the PX domain of p47phox to phosphatidylinositol 3, 4-bisphosphate and phosphatidic acid is masked by an intramolecular interaction. The EMBO journal.

13. Chandra, M., Y. K.-Y. Chin, C. Mas, J. R. Feathers, B. Paul, S. Datta, K.-E. Chen, X. Jia, Z. Yang, S. J. Norwood, et al., 2019. Classification of the human phox homology (PX) domains based on their phosphoinositide binding specificities. Nature communications 10:1528.

14. Zhang, Y., X. Pang, J. Li, J. Xu, V. W. Hsu, and F. Sun, 2021. Structural insights into membrane remodeling by SNX1. Proceedings of the National Academy of Sciences 118:e2022614118.

15. Simonetti, B., C. M. Danson, K. J. Heesom, and P. J. Cullen, 2017. Sequence-dependent cargo recognition by SNX-BARs mediates retromer-independent transport of CI-MPR. Journal of Cell Biology 216:3695–3712.

16. Kvainickas, A., A. Jimenez-Orgaz, H. Nägele, Z. Hu, J. Dengjel, and F. Steinberg, 2017. Cargo-selective SNX-BAR proteins mediate retromer trimer independent retrograde transport. Journal of Cell Biology 216:3677–3693.

17. Yong, X., L. Zhao, W. Deng, H. Sun, X. Zhou, L. Mao, W. Hu, X. Shen, Q. Sun, D. D. Billadeau, et al., 2020. Mechanism of cargo recognition by retromer-linked SNX-BAR proteins. PLoS biology 18:e3000631.

18. Simonetti, B., B. Paul, K. Chaudhari, S. Weeratunga, F. Steinberg, M. Gorla, K. J. Heesom, G. J. Bashaw, B. M. Collins, and P. J. Cullen, 2019. Molecular identification of a BAR domain-containing coat complex for endosomal recycling of transmembrane proteins. Nature cell biology 21:1219–1233.

19. Shortill, S. P., M. S. Frier, and E. Conibear, 2022. You can go your own way: SNX-BAR coat complexes direct traffic at late endosomes. Current Opinion in Cell Biology 76:102087.

20. Koharudin, L. M., W. Furey, H. Liu, Y.-J. Liu, and A. M. Gronenborn, 2009. The Phox Domain of Sorting Nexin 5 Lacks Phosphatidylinositol 3-Phosphate (PtdIns (3) P) Specificity and Preferentially Binds to Phosphatidylinositol 4, 5-Bisphosphate (PtdIns (4, 5) P2)*. Journal of Biological Chemistry 284:23697–23707.

21. Simonetti, B., C. M. Danson, K. J. Heesom, and P. J. Cullen, 2017. Sequence-dependent cargo recognition by SNX-BARs mediates retromer-independent transport of CI-MPR. Journal of Cell Biology 216:3695–3712.

22. Lopez-Robles, C., S. Scaramuzza, E. N. Astorga-Simon, M. Ishida, C. D. Williamson, S. Baños-Mateos, D. Gil-Carton, M. Romero-Durana, A. Vidaurrazaga, J. Fernandez-Recio, et al., 2023. Architecture of the ESCPE-1 membrane coat. Nature Structural & Molecular Biology 30:958–969.

23. Ramakrishnan, N., P. Sunil Kumar, and J. H. Ipsen, 2010. Monte Carlo simulations of fluid vesicles with in-plane orientational ordering. Physical Review E—Statistical, Nonlinear, and Soft Matter Physics 81:041922.

24. Ramakrishnan, N., P. S. Kumar, and R. Radhakrishnan, 2014. Mesoscale computational studies of membrane bilayer remodeling by curvature-inducing proteins. Physics reports 543:1–60.

25. Bonazzi, F., C. K. Hall, and T. R. Weikl, 2021. Membrane morphologies induced by mixtures of arc-shaped particles with opposite curvature. Soft Matter 17:268–275.

26. Duncan, A. L., and W. Pezeshkian, 2023. Mesoscale simulations: an indispensable approach to understand biomembranes. Biophysical Journal 122:1883–1889.

27. Kumar, G., and A. Srivastava, 2022. Membrane remodeling due to a mixture of multiple types of curvature proteins. Journal of Chemical Theory and Computation 18:5659–5671.

28. Kumar, G., S. C. Duggisetty, and A. Srivastava, 2022. A review of mechanics-based mesoscopic membrane remodeling methods: capturing both the physics and the chemical diversity. The Journal of Membrane Biology 255:757–777.

29. Zhong, Q., M. J. Watson, C. S. Lazar, A. M. Hounslow, J. P. Waltho, and G. N. Gill, 2005. Determinants of the endosomal localization of sorting nexin 1. Molecular biology of the cell 16:2049–2057.

30. Natarajan, C., and A. Srivastava, 2024. Efficiently determining membrane-bound conformations of peripheral membrane proteins using replica exchange with hybrid tempering. Eur. Phys. J. Spec. Top. 3039–3051.

31. Appadurai, R., J. Nagesh, and A. Srivastava, 2021. High resolution ensemble description of metamorphic and intrinsically disordered proteins using an efficient hybrid parallel tempering scheme. Nature communications 12:1–11.

32. Jo, S., T. Kim, V. G. Iyer, and W. Im, 2008. CHARMM-GUI: a web-based graphical user interface for CHARMM. Journal of computational chemistry 29:1859–1865.

33. Van Der Spoel, D., E. Lindahl, B. Hess, G. Groenhof, A. E. Mark, and H. J. Berendsen, 2005. GROMACS: fast, flexible, and free. J. Comp. Chem 26:1701–1718.

34. Nosé, S., 1984. A unified formulation of the constant temperature molecular dynamics methods. J. Chem. Phys 81:511–519.

35. Hoover, W. G., 1985. Canonical dynamics: Equilibrium phase-space distributions. Phys. Rev. A 31:1695.

36. Parrinello, M., and A. Rahman, 1981. Polymorphic transitions in single crystals: A new molecular dynamics method. J. App. Phys 52:7182–7190.

37. Bauer, P., B. Hess, and E. Lindahl, 2023. GROMACS 2022.5 Source Code. Zenodo.

38. The PLUMED consortium, 2019. Promoting transparency and reproducibility in enhanced molecular simulations. Nature Methods 16:670–673. 10.1038/s41592-019-0506-8.

39. Huang, J., S. Rauscher, G. Nawrocki, T. Ran, M. Feig, B. L. De Groot, H. Grubmüller, and A. D. MacKerell Jr, 2017. CHARMM36m: an improved force field for folded and intrinsically disordered proteins. Nature methods 14:71–73.

40. Jorgensen, W. L., J. Chandrasekhar, J. D. Madura, R. W. Impey, and M. L. Klein, 1983. Comparison of simple potential functions for simulating liquid water. The Journal of chemical physics 79:926–935.

41. Natarajan, C., and A. Srivastava, 2024. Efficiently determining membrane-bound conformations of peripheral membrane proteins using replica exchange with hybrid tempering. The European Physical Journal Special Topics 233:3039–3051.

42. Yang, J., R. Yan, A. Roy, D. Xu, J. Poisson, and Y. Zhang, 2015. The I-TASSER Suite: protein structure and function prediction. Nature methods 12:7–8.

43. Zhang, Y., 2008. I-TASSER server for protein 3D structure prediction. BMC bioinformatics 9:1–8.

44. Lopéz-Blanco, J. R., and P. Chacón, 2013. iMODFIT: efficient and robust flexible fitting based on vibrational analysis in internal coordinates. Journal of structural biology 184:261–270.

45. Pettersen, E. F., T. D. Goddard, C. C. Huang, G. S. Couch, D. M. Greenblatt, E. C. Meng, and T. E. Ferrin, 2004. UCSF Chimera—a visualization system for exploratory research and analysis. Journal of computational chemistry 25:1605–1612.

46. Scheurer, M., P. Rodenkirch, M. Siggel, R. C. Bernardi, K. Schulten, E. Tajkhorshid, and T. Rudack, 2018. PyContact: rapid, customizable, and visual analysis of noncovalent interactions in MD simulations. Biophysical journal 114:577–583.

47. Canham, P. B., 1970. The minimum energy of bending as a possible explanation of the biconcave shape of the human red blood cell. Journal of theoretical biology 26:61–81.

48. Helfrich, W., 1973. Elastic properties of lipid bilayers: theory and possible experiments. Zeitschrift für Naturforschung c 28:693–703.

49. Ramakrishnan, N., P. S. Kumar, and J. H. Ipsen, 2010. Monte Carlo simulations of fluid vesicles with in-plane orientational ordering. Physical Review E 81:041922.

50. Lebwohl, P., 1972. PA Lebwohl and G. Lasher, Phys. Rev. A 6, 426 (1972). Phys. Rev. A 6:426.

51. Nakazawa, K., G. Kumar, B. Chauvin, A. Di Cicco, L. Pellegrino, M. Trichet, B. Hajj, J. Cabral, A. Sain, S. Mangenot, et al., 2023. A human septin octamer complex sensitive to membrane curvature drives membrane deformation with a specific mesh-like organization. Journal of Cell Science 136.

52. Kumar, G., N. Ramakrishnan, and A. Sain, 2019. Tubulation pattern of membrane vesicles coated with biofilaments. Physical Review E 99:022414.

53. Sreeja, K., J. H. Ipsen, and P. S. Kumar, 2015. Monte Carlo simulations of fluid vesicles. Journal of Physics: Condensed Matter 27:273104.

54. Ye, B., W. Tian, B. Wang, and J. Liang, 2024. CASTpFold: Computed Atlas of Surface Topography of the universe of protein Folds. Nucleic Acids Research 52:W194–W199.

55. Overduin, M., and R. Bhat, 2024. Recognition and remodeling of endosomal zones by sorting nexins. Biochimica et Biophysica Acta (BBA)-Biomembranes 184305.

56. Veretenenko, I. I., Y. A. Trofimov, N. A. Krylov, and R. G. Efremov, 2024. Nanoscale lipid domains determine the dynamic molecular portraits of mixed DOPC/DOPS bilayers in a fluid phase: A computational insight. Biochimica et Biophysica Acta (BBA) - Biomembranes 1866:184376.

57. Corradi, V., E. Mendez-Villuendas, H. I. Ingólfsson, R.-X. Gu, I. Siuda, M. N. Melo, A. Moussatova, L. J. DeGagné, B. I. Sejdiu, G. Singh, T. A. Wassenaar, K. Delgado Magnero, S. J. Marrink, and D. P. Tieleman, 2018. Lipid–Protein Interactions Are Unique Fingerprints for Membrane Proteins. ACS Central Science 4:709–717.

58. Dowhan, W., 2017. Understanding phospholipid function: Why are there so many lipids? Journal of Biological Chemistry 292:10755–10766.

59. Kervin, T., and M. Overduin, 2024. Membranes are functionalized by a proteolipid code. BMC Biology 22.

60. Soteriou, C., M. Xu, S. D. Connell, A. I. Tyler, A. C. Kalli, and J. L. Thorne, 2025. Two cooperative lipid binding sites within the pleckstrin homology domain are necessary for AKT binding and stabilization to the plasma membrane. Structure 10.1016/j.str.2024.10.020.

61. Krick, R., R. A. Busse, A. Scacioc, M. Stephan, A. Janshoff, M. Thumm, and K. Kühnel, 2012. Structural and functional characterization of the two phosphoinositide binding sites of PROPPINs, a - propeller protein family. Proceedings of the National Academy of Sciences 109:E2042–E2049. https://www.pnas.org/doi/abs/10.1073/pnas.1205128109.

62. Srivastava, A., 2025. It takes two to tango: The second membrane-binding site in peripheral proteins. Structure 33:10–12.

